# Cohesin promotes stochastic domain intermingling to ensure proper regulation of boundary-proximal genes

**DOI:** 10.1101/649335

**Authors:** Jennifer M. Luppino, Daniel S. Park, Son C. Nguyen, Yemin Lan, Zhuxuan Xu, Eric F. Joyce

**Affiliations:** Department of Genetics, Penn Epigenetics Institute, Perelman School of Medicine, University of Pennsylvania, Philadelphia, PA 19104; Department of Cell and Developmental Biology, Penn Epigenetics Institute, Perelman School of Medicine, University of Pennsylvania, Philadelphia, PA 19104, USA.

## Abstract

The mammalian genome can be segmented into thousands of topologically associated domains (TADs) based on chromosome conformation capture studies, such as Hi-C. TADs have been proposed to act as insulated neighborhoods, spatially sequestering and insulating the enclosed genes and regulatory elements through chromatin looping and self-association. Recent results indicate that inter-TAD interactions can also occur, suggesting boundaries may be semi-permissible. However, the nature, extent, and function, if any, of these inter-TAD interactions remains unclear. Here, we combine super-and high-resolution microscopy with Oligopaint technology to precisely quantify the interaction frequency within and between neighboring domains in human cells. We find that intermingling across domain boundaries is a widespread feature of the human genome, with varying levels of interactions across different loci that correlate with their differing boundary strengths by Hi-C. Moreover, we find that cohesin depletion, which is known to abolish TADs at the population-average level, does not induce ectopic interactions but instead reduces both intra- and inter-domain interactions to a similar extent. Reduced chromatin intermixing due to cohesin loss affects domain incorporation and transcriptional bursting frequencies of genes close to architectural boundaries, potentially explaining the gene expression changes observed in the cohesinopathy Cornelia de Lange syndrome. Together, our results provide a mechanistic explanation for stochastic domain intermingling, arguing that cohesin partially bypasses boundaries to promote alternating incorporation of boundary-proximal genes into neighboring regulatory domains.

## Introduction

Chromosomes are folded at different length scales in the nucleus of eukaryotic cells (Bickmore, 2013; Gibcus and Dekker, 2013). At the largest scale, chromosomes are spatially partitioned away from each other into chromosome territories (Cremer and Cremer, 2010). Chromosome conformation capture-based methods, including Hi-C, have further subdivided the genome into topologically associated domains (TADs), nested sub-TADs, and discrete chromatin loops (Dixon et al., 2012; Hansen et al., 2017; Kim et al., 2018; Nora et al., 2012; Phillips-Cremins et al., 2013; Rao et al., 2014; Rao et al., 2017). This level of genome organization has been proposed to regulate many cellular processes, including gene expression, DNA replication, and DNA damage repair, though the exact mechanisms by which chromatin structure facilitates these functions remain unclear.

Several architects of genome organization have been identified. In particular, the insulator protein CTCF and ring-shaped cohesin complex colocalize on chromatin (Wendt et al., 2008) at the anchors of loops (Rao et al., 2014; Splinter et al., 2006) and the boundaries of TADs (Dixon et al., 2012; Lieberman-Aiden et al., 2009; Nora et al., 2012; Rao et al., 2014). This has led to the hypothesis that CTCF dimerization halts cohesin-mediated chromatin extrusion to form the boundaries of contact domains (Goloborodko et al., 2016; Mirny et al., 2019; Sanborn et al., 2015). Consistent with a role for these factors in genome organization, depletion of either CTCF or cohesin from chromatin greatly perturbs loop and TAD formation at the population average level and induces widespread, yet moderate effects on gene expression (Nora et al., 2017; Rao et al., 2017; Schwarzer et al., 2017; Wutz et al., 2017). More dramatically, deletion of CTCF binding sites at the boundary of select TADs can result in ectopic transcriptional activation of one or more flanking genes via formation of an enhancer-promoter loop across the deleted boundary domain (Dowen et al., 2014; Lupianez et al., 2015). Thus, TADs and sub-TADs are presumed to constrain gene regulation within their boundaries (Ji et al., 2016; Sun et al., 2019), thereby simultaneously insulating and promoting the specificity or fidelity of an enhancer for its target gene (Levine et al., 2014; Sun et al., 2019). Importantly, insulation is believed to function via the spatial separation of TADs from each other. However, recent single-cell Hi-C datasets and imaging-based approaches have suggested extensive heterogeneity in both TAD and sub-TAD organization at the single-cell level (Bintu et al., 2018; Finn et al., 2019; Flyamer et al., 2017; Nagano et al., 2013). Yet, the molecular determinants of these heterogeneous interactions and their relevance to gene expression remain unknown.

To address this issue, we generated Oligopaint-based probes for fluorescence in situ hybridization (FISH) (Beliveau et al., 2015; Beliveau et al., 2012; Beliveau et al., 2018) to precisely target entire Hi-C-defined TADs in single cells. We use a combination of high- and super-resolution microscopy to quantify the frequency and extent of spatial intermingling between neighboring domains of different length scales and chromatin types. These approaches reveal extensive interactions between neighboring domains that are promoted by the cohesin complex. Reduced chromatin intermixing due to cohesin loss affects domain incorporation and transcriptional bursting frequencies of genes close to architectural boundaries. Therefore, we propose that, rather than forming spatially insulated domains, cohesin promotes stochastic domain intermingling to ensure proper regulation of boundary-proximal genes.

## Results

### Cell-to-cell variability of inter-TAD association is a widespread feature of the human genome

To measure the frequency and extent of interactions between entire TADs at the single chromosome level, we designed a FISH-based assay that tiles probes across population-based contact domains using the programmable Oligopaint technology (Beliveau et al., 2014; Beliveau et al., 2015; Beliveau et al., 2017; Beliveau et al., 2012; Beliveau et al., 2018). We then applied our “TAD FISH” assay in HCT-116, a human colorectal carcinoma cell line from which high-resolution Hi-C maps were previously generated (Rao et al., 2017).

We determined TAD and sub-TAD coordinates using the TopDom TAD caller (Shin et al., 2016) from previously published in situ Hi-C data (Rao et al., 2017), identifying 4,429 contact domains with a median size of 425 kb (**Table S1**). The boundaries of contact domains were further refined by co-localization of CTCF and RAD21, based on ENCODE ChIP-Seq datasets (Consortium, 2012). We then designed Oligopaint probes targeting two pairs of consecutive TADs located on chromosomes 12 and 22 (**Table S2**). These TAD pairs were chosen based on their relative TAD boundary strength (20^th^ and 80^th^ percentile, respectively) as well as their differing levels of cohesin, gene density, expression status, and chromatin modifications (**Figures 1A and S1**). In particular, the shared boundary between the TAD pair on chromosome 12 is contained within an active A compartment whereas the TAD pair on chromosome 22 is mostly contained within a silent B compartment. We performed two-color FISH to alternatively label neighboring TADs in a pair-wise fashion.

**Figure 1.**
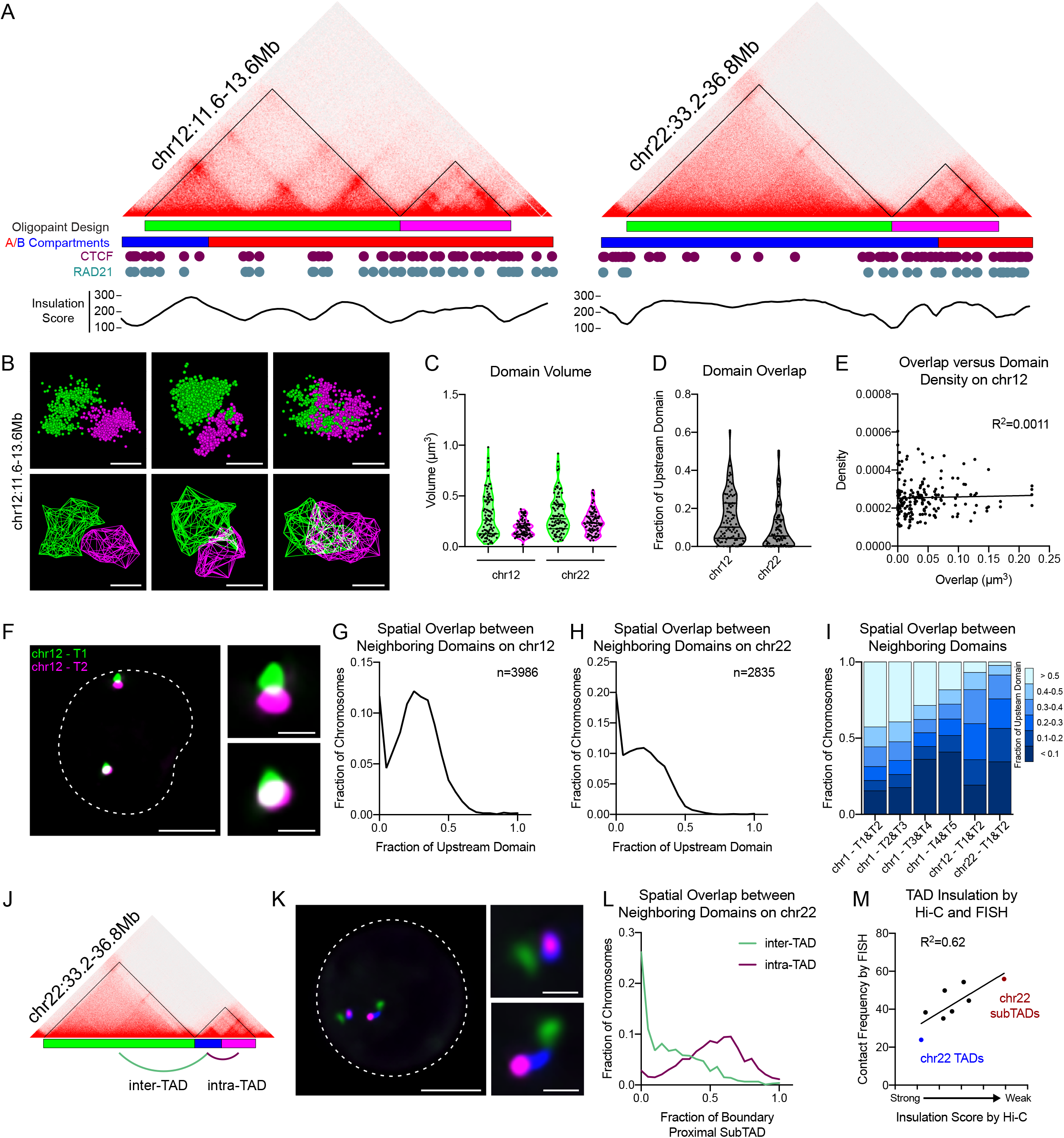
Cell-to-cell variability of inter-TAD association is a widespread feature of the human genome. (A) Hi-C contact matrices and Oligopaint probe design for neighboring TADs on chromosomes 12 (left) and 22 (right) in the HCT-116 cell line, with corresponding A/B compartment designations, CTCF and RAD21 binding profiles, and Hi-C insulation scores based on mean contact frequency at 25 kb bins. (B) Three representative 3D-STORM images of neighboring TADs on chr12:11.6Mb-13.6Mb illustrated by individual fluorescence localizations (above) and 3D hull reconstruction (below). Scale bar equals 500 nm. (C) Violin plots of TAD volume quantified from 3D-STORM images. n > 90 chromosomes. (D) Violin plots of spatial overlap between neighboring domains, normalized to the volume of the upstream domain. n > 90 chromosomes. (E) Scatterplot of the volume of spatial overlap between neighboring domains at chr12:11.6Mb-13.6Mb (x-axis) versus the particle density of either domain (y-axis). n = 91 chromosomes. (F) Representative deconvolved widefield image of neighboring TADs at chr12:11.Mb-13.6Mb. Dashed lines represent nuclear edges. Scale bar equals 5 μm (left) or 1 μm (zoomed images, right). (G) Histogram of spatial overlap between neighboring domains on chr12:11.6Mb-13.6Mb, normalized to the volume of the upstream domain. n = 3986 chromosomes. (H) Histogram of spatial overlap between neighboring domains on chr22:33.2Mb-36.8Mb, normalized to the volume of the upstream domain. n = 2835 chromosomes. (I) Stacked bar graph of spatial overlap between six pairs of neighboring TADs. Overlap presented as a fraction of upstream domain volume. n > 2294 chromosomes. (J) Hi-C contact matrix of chr22:33.2Mb-36.8Mb and Oligopaint design corresponding to (K-L). (K) Representative deconvolved widefield image of three-color FISH to chr22:33.2Mb-36.8Mb. Dashed lines represent nuclear edges. Scale bar equals 5 μm (left) or 1 μm (zoomed images, right). (L) Histograms of inter-TAD overlap (green) and intra-TAD overlap (purple). Overlap normalized to the volume of the boundary-proximal subTAD. n = 1060 chromosomes. (M) Scatterplot of boundary insulation score determined by TopDom analysis of Hi-C versus the contact frequencies of neighboring domains by FISH. Each point represents the average of two biological replicates for eight neighboring domain pairs.

We first applied super-resolution microscopy using 3D stochastic optical reconstruction (3D-STORM) to measure the volume and spatial overlap between adjacent domains with <50 nm error in their localization and <5% error in their physical sizes (Beliveau et al., 2015). Cells were synchronized in G1 to avoid heterogeneity due to the cell cycle or presence of sister chromatids (**Figure S2A**). We observed a wide diversity in the shape, volume, and particle density of each reconstructed domain across the cell population (**Figures 1B-C and S3A**). While alleles within the same cell showed a moderate correlation in domain volume, no such correlation was observed between neighboring domains across the same chromosome, indicating stochastic compaction rates were intrinsic to each chromatin domain (**Figures S3B-E**).

To quantify interactions between neighboring domains, we measured the inter-TAD overlap volume, presented as a fraction of domain volume (**Figure 1D**). If TADs exist as spatially separate structures, we would expect little to no overlap between the reconstructed domains. This was indeed the case for 15-34% of chromosomes depending on the locus. However, the majority of chromosomes (66-85%) showed some level of intermingling between the probed domain pairs. Neighboring domains overlapped by up to 50-61%, indicating a high level of variability in inter-domain interactions within the cell population. The amount of spatial overlap between neighboring domains, however, did not correlate with the volume or particle density of either domain; therefore, interactions between the domains did not seem to be a direct result of intra-domain decompaction (**Figures 1E and S3F-H**).

To draw a distribution of inter-domain interactions across the cell population, we next applied diffraction-limited microscopy and captured a minimum of 2,000 chromosomes per domain pair (**Figure 1F**). Images were subjected to maximum likelihood deconvolution followed by custom 3D segmentation, which allowed us to trace the 3D edges of each TAD signal and plot the distribution of their volumes and spatial overlap based on voxel colocalization (**Figure S3I**). The spatial overlap between neighboring domains recapitulated the results observed by super-resolution microscopy; only 12-20% of neighboring domains were spatially separated (**Figures 1G-H**). Interestingly, two populations emerged from the distributions of spatial overlap at both loci, with one of the largest peaks consistently at little to no overlap and the other representing a wide range of overlap fractions (**Figures 1G-H**). We tested an additional four TAD pairs on chromosome 1 and observed similar patterns of inter-TAD associations, the extent of which differed in a locus-specific manner (**Figures 1I, S1, and S3J-O**). Similar results were also observed in asynchronous cell populations, indicating this is not a feature specific to cells in G1 (**Figure S3P**).

To compare inter-TAD to intra-TAD interactions, we separately labeled two subdomains within the chromosome 22 TAD locus and measured their spatial overlap (**Figure 1J-L**). Higher overlap fractions were frequently observed between intra-TAD domains (**Figure 1L**), consistent with greater levels of intra-TAD interactions as predicted by Hi-C. Surprisingly, no correlation was observed (R^2^=0.095) between intra- and inter-TAD overlap on the same chromosome (**Figure S3Q**), suggesting that stochastic interactions across intra- and inter-TAD boundaries occur independently. To compare our FISH data to that of Hi-C, we plotted the frequency of domain interactions using a voxel volume cutoff (500 nm^3^) to the insulation score of their shared boundary based on the mean interaction frequency with 25kb bins. Combining data from all intra- and inter-TAD pairs tested, we find a strong correlation between these two metrics (R^2^=0.62; **Figure 1M**), suggesting Hi-C and our FISH assay are in agreement when comparing relative contact frequencies across different boundaries. However, our data further suggest that extensive interactions between neighboring domains are frequent events, even across the strongest boundaries in the human genome.

### Interactions between contact domains predominantly occur across the boundary

To determine the nature of the interactions between neighboring TADs, we first divided the upstream chromosome 12 TAD into its three subdomains (S1, S2, and S3) and generated Oligopaint probes targeting each (**Figures 2A**). Consistent with high levels of intra-TAD associations, the subdomains contacted each other in 80-92% of cells and frequently displayed >50% overlap fractions (**Figures 2B-D**). We then measured the spatial overlap of each subdomain with the downstream TAD (T2) to determine if inter-TAD association is largely dependent on distance or more complex 3D interactions (**Figures 2E-F**). Interestingly, all three subdomains (S1, S2, S3) came into contact with the neighboring TAD in ≥78% of cells (**Figure 2G**). However, the boundary-proximal subdomain (S3) exhibited the most overlap with the downstream TAD (**Figure 2H**). These data show that, while all parts of the upstream TAD can come into contact with the downstream TAD, inter-TAD interactions are more frequently intermixed with boundary-proximal chromatin.

**Figure 2.**
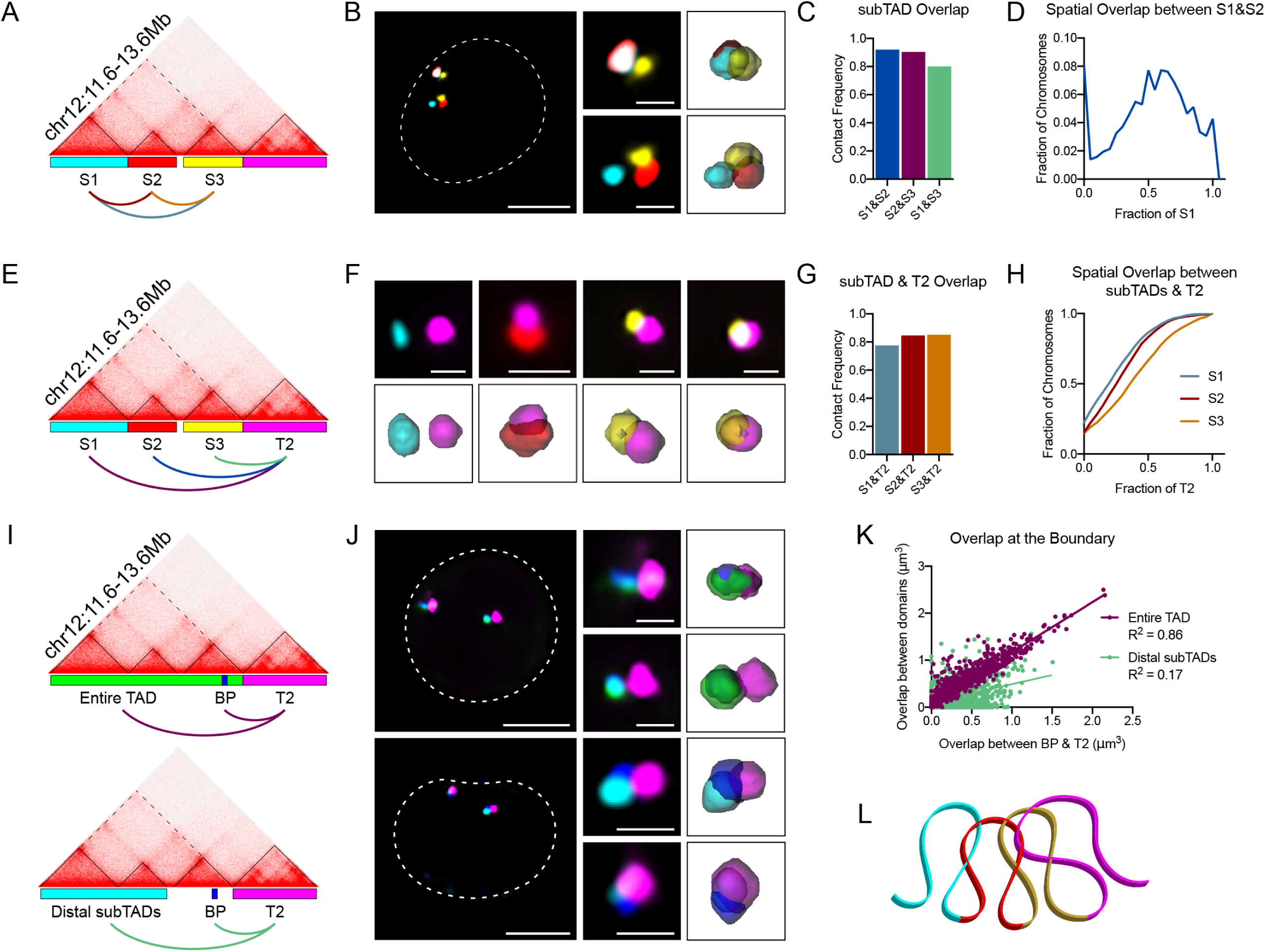
Interactions between contact domains predominantly occur across the boundary. (A) Hi-C contact matrix of chr12:11.6Mb-13.6Mb and Oligopaint design corresponding to (B-D). (B) Representative deconvolved widefield image of three-color FISH experiments illustrating interactions between the three subTADs. Dashed white lines represent nuclear edges. Scale bar equals 5 μm (left) or 1 μm (zoomed images, right). Corresponding 3D segmentation of FISH signals on right. (C) Fraction of chromosomes with contact between subTADs. n > 1063 chromosomes. (D) Histogram of spatial overlap between subTADs S1 and S2, normalized to the volume of S1. n = 1076 chromosomes. (E) Hi-C contact matrix of chr12:11.6Mb-13.6Mb and Oligopaint design corresponding to (F-H). (F) Representative deconvolved widefield image of FISH experiments illustrating interactions between the three subTADs and the downstream domain T2. Scale bar equals 1 μm. Corresponding 3D segmentation of FISH signals below each image. (G) Fraction of chromosomes with contact between subTADs and downstream TAD (T2). n > 1932 chromosomes. (H) Cumulative distribution plot of spatial overlap between each subTAD with the downstream TAD (T2), normalized to the volume of T2. n > 1932 chromosomes. (I) Hi-C contact matrix and Oligopaint design corresponding to (J-K). “BP” represents the boundary-proximal region. (J) Representative deconvolved widefield images of three-color FISH corresponding to (I). Dashed white lines represent nuclear edges. Scale bar equals 5 μm (left) or 1 μm (zoomed images, middle). Corresponding 3D segmentation of FISH signals on right. (K) Scatterplot of spatial overlap between BP and T2 (x-axis) and either the entire TAD (purple) or distal subTADs (green). n > 1005 chromosomes. (L) Cartoon schematic of chr12:11.6Mb-13.6Mb locus representing both local, distance-dependent looping interactions between neighboring sub-TADs and more complex, distal 3D interactions between S2 (blue) and T2 (purple).

Next, we designed probes targeting a 33 kb boundary-proximal region between the TADs and performed three-color FISH to additionally label the neighboring TADs (**Figures 2I-J**). We found a strong linear relationship (R^2^=0.86) between the spatial overlap of the boundary-proximal region and downstream TAD with the overlap between neighboring TADs (**Figure 2K**). Therefore, the extent of overlap between neighboring TADs is strongly associated with the boundary proximal region. However, this was not true when examining the two distal subdomains only. Their collective overlap with the downstream TAD only weakly correlated with overlap between the boundary-proximal domain and downstream TAD (R^2^=0.17) (**Figure 2K**). This suggests that when interactions between distal portions of TADs do occur, they often do not include boundary-proximal chromatin. Similar results were found at an additional locus on chromosome 22 (**Figures S4A-C**). Together, these data suggest that, while more complex 3D interactions can contribute to the variable interactions between neighboring domains, the majority of their intermixing reflect their genomic distance and therefore occur near the boundary (**Figure 2L**).

### Cohesin promotes interactions across domain boundaries

To test the contribution of cohesin to inter-domain interactions, we carried out an acute depletion of RAD21, a core component of the cohesin ring, via auxin-inducible degradation (AID) in HCT-116 cells (Natsume et al., 2016). Hi-C in this cell line has previously shown complete loss of loop domains following RAD21 degradation (**Figures 3A-B**) (Rao et al., 2017). Our reanalysis of these data using TopDom TAD detection (Shin et al., 2016) revealed that 75% of TAD boundaries were lost following auxin treatment (**Table S3**). To determine if this loss was associated with ectopic interactions across boundaries, we repeated our FISH assay in synchronized and arrested G1 cells treated with auxin for 6 hours. This treatment resulted in a >95% reduction in chromatin-bound RAD21 levels (**Figures S2B-D**).

**Figure 3.**
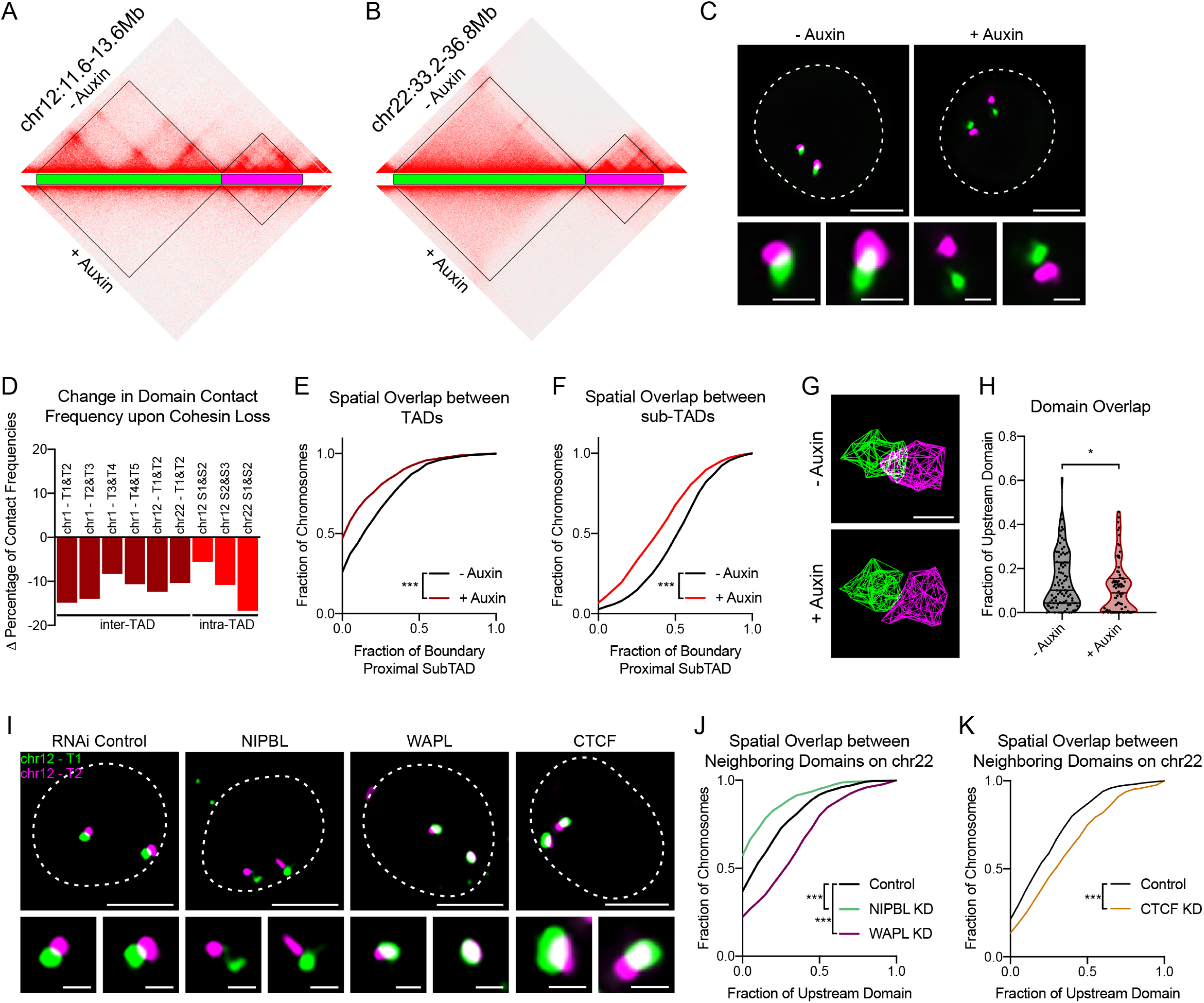
Cohesin promotes interactions across domain boundaries. (A) Hi-C contact matrix of chr12:11.6Mb-13.6Mb in untreated and auxin treated HCT-116-RAD21-AID cells. (B) Hi-C contact matrix of chr22:33.2-36.8Mb in untreated and auxin treated HCT-116-RAD21-AID cells. (C) Representative deconvolved widefield images of neighboring TADs across chr12:11.6Mb-13.6Mb with or without auxin treatment. Dashed lines represent nuclear edges. Scale bar equals 5 μm (left) or 1 μm (zoomed images, below). (D) Difference in percent contact frequencies between domains following auxin treatment. Each bar represents average of two biological replicates. (E) Cumulative distribution plot of spatial overlap between TADs on chr22:33.2-36.8Mb, normalized to the volume of T2. *** p < 0.001, two-tailed Mann Whitney. n > 1060 chromosomes. (F) Cumulative distribution plot of spatial overlap between subTADs on chr22:33.2-36.8Mb, normalized to the volume of T2. *** p < 0.001, two-tailed Mann Whitney. n > 1060 chromosomes. (G) Representative 3D-STORM images of neighboring TADs on chr22:33.2-36.8Mb. Scale bar equals 500 nm. (H) Violin plot of spatial overlap between domains on chr22:33.2-36.8Mb, normalized to volume of the upstream domain. * p = 0.044, two-tailed Mann Whitney test. n > 76 chromosomes. (I) Representative deconvolved widefield images of neighboring TADs on chr12:11.6Mb-13.6Mb in RNAi control, NIPBL, WAPL, and CTCF depleted HCT-116 cells. Dashed lines represent nuclear edges. Scale bar equals 5 μm (left) or 1 μm (zoomed images, below). (J) Cumulative frequency distribution of spatial overlap between neighboring domains on chr22:33.2-36.8Mb in control, NIPBL, and WAPL depleted cells. Overlap normalized to the volume of the upstream domain. *** p < 0.001, two-tailed Mann Whitney test. n > 636 chromosomes. (K) Cumulative frequency distribution of spatial overlap between neighboring domains on chr22:33.2Mb-36.8Mb in control and CTCF depleted cells. Overlap normalized to the volume of the upstream domain. *** p < 0.001, two-tailed Mann Whitney test. n > 332 chromosomes.

Despite the loss of boundaries by Hi-C, all nine domain pairs tested exhibited reduced contact frequencies following RAD21 degradation (**Figure 3C-D**). For example, contact between chromosome 22 TAD pairs was decreased by 10%, whereas intra-TAD domains at the same locus was reduced by 17% (**Figure 3D**). Although cohesin loss affected some domain pairs more than others, these locus-specific differences did not seem to reflect their boundary strength prior to treatment or the size or type of domain pair being tested (**Figure S5A**). Furthermore, when overlap was observed, the amount shifted towards significantly less intermixing following RAD21 degradation (p<0.001; **Figures 3E-F and S5B-I**). As in control cells, no correlation between intra- and inter-TAD overlap was observed (**Figure S5J**). We validated these findings using 3D-STORM, which revealed an increase in spatial separation between domains following cohesin loss (**Figures 3G-H and S5K-N**). Surprisingly, the volume and localization density of each domain did not increase following cohesin loss (**Figure S5O-P**). Instead, local compaction rates remained mostly unchanged, with a few domains becoming slightly more compact following auxin treatment. Taken together, this suggests that, while cohesin loss may not alter local chromatin compaction, higher-order interactions between intra- and inter-TADs decrease overall.

### WAPL and CTCF restrict cohesin-dependent interactions across domain boundaries

To further validate our results, we next sought to determine if the regulation of cohesin would also affect the level of spatial overlap between neighboring domains. Cohesin is loaded onto chromatin by NIPBL, whereas the negative regulator of cohesin, WAPL, opens the cohesin ring to release it from DNA (Nasmyth and Haering, 2009). We depleted NIPBL and WAPL using RNAi in HCT-116 cells to decrease and increase cohesin occupancy, respectively, and then measured the overlap between neighboring domains at two loci by FISH (**Figures 3I-J and S2E**). NIPBL-depleted chromatin showed significantly (p<0.001) less contact and more spatial separation between domains, similar to RAD21 degradation, whereas WAPL-depleted chromatin showed significantly (p<0.001) more contact and greater overlap between the domains (**Figures 3J and S5Q**). Double knockdown of NIPBL and WAPL phenocopied NIPBL depletion alone, indicating the increase in overlap was cohesin-dependent (**Figures S5R-T**). Finally, we depleted the insulator protein CTCF, which similar to cohesin, perturbs TADs at the population-average level (Nora et al., 2017). In contrast to cohesin loss, however, we observed significantly increased contact and spatial overlap between neighboring TADs (p<0.001), similar to WAPL depletion (**Figure 3K**). This indicates that depletion of cohesin and CTCF show opposite effects on inter-TAD interactions. Taken together, these results indicate that CTCF and WAPL restrict but do not eliminate cohesin-dependent interactions across TAD boundaries.

### Expression of boundary-proximal genes is sensitive to cohesin levels

TADs have long been implicated in the regulation of gene expression and previous work using nascent RNA-sequencing revealed 4,196 genes to be significantly differentially expressed in HCT-116 cells following RAD21 degradation, albeit with relatively small changes (Rao et al., 2017). Of 1,607 genes showing a >30% change in expression, only 64 changed by >2-fold (**Figures 4A-B**). Given that domains were less likely to interact in the absence of cohesin, we reexamined the position of these genes in the context of domain boundaries.

**Figure 4.**
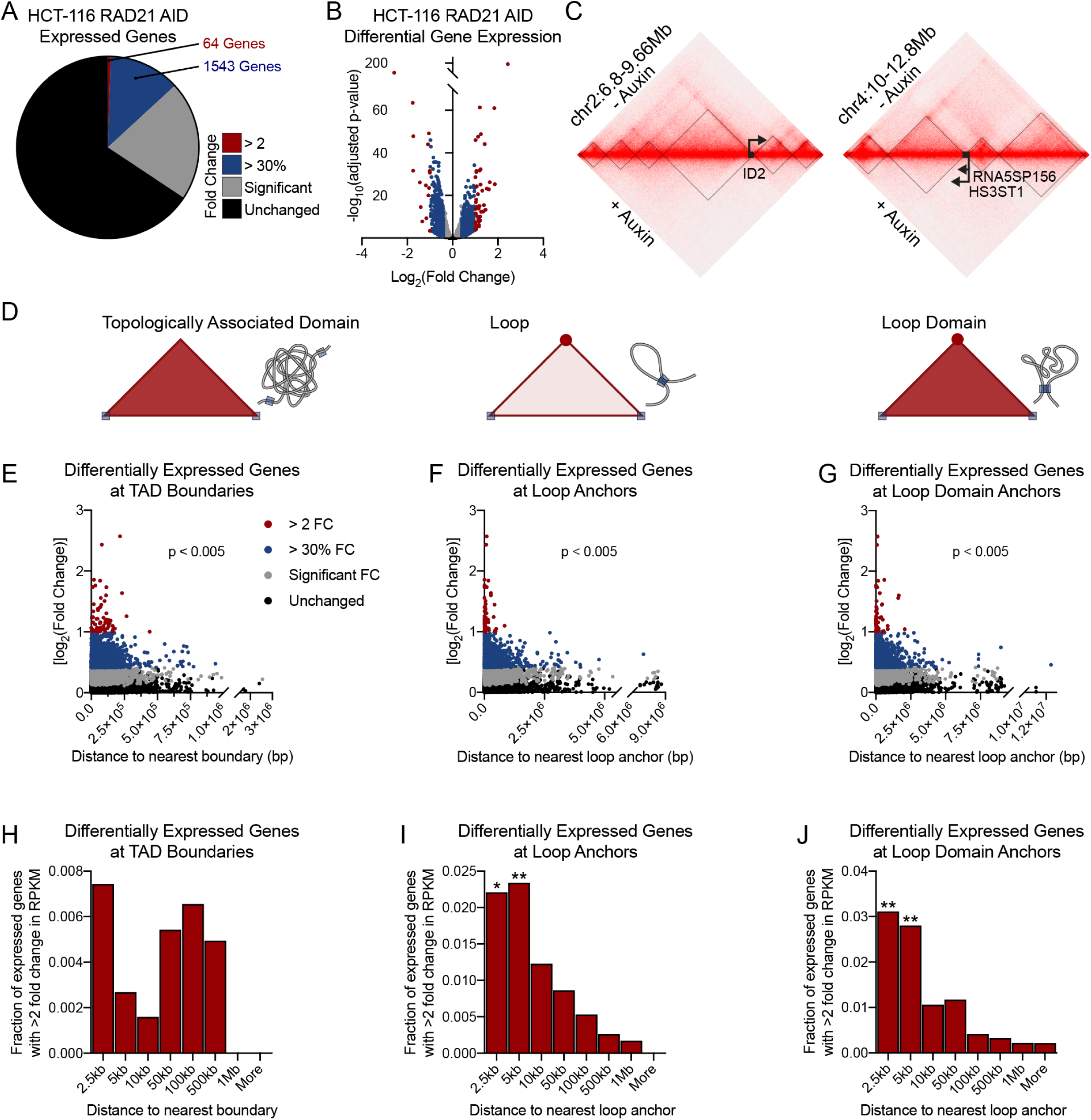
Expression of boundary-proximal genes is sensitive to cohesin levels. (A) Pie chart representing genes in HCT116-RAD21-AID cells that are differentially expressed upon auxin treatment by PRO-Seq data published in Rao, *et al*. (Rao et al., 2017). (B) Volcano plot of the log_2_(fold change) (x-axis) versus significance (y-axis) of differentially expressed genes (DEGs) by PRO-Seq data published in Rao, *et al*. (Rao et al., 2017). (C) Hi-C contact matrices for two regions depicting three differentially expressed genes (DEGs) (ID2, RNA5SP156, and HS3ST1) near TAD boundaries. Black lines represent TopDom TAD calls. Arrows represent relative gene locations and directionality. (D) Cartoons depicting Hi-C contact matrices and predicted chromatin folding of TADs, loops, and loop domains. Boundaries/anchors depicted as blue boxes. (E) Scatterplot of absolute log_2_(fold change) of DEGs versus the distance between their TSS and the nearest TAD boundary. p = 0.0043, Spearman correlation. (F) Scatterplot of absolute log_2_(fold change) of DEGs versus the distance between their TSS and the nearest loop anchor. p = 0.0001, Spearman correlation. (G) Scatterplot of absolute log_2_(fold change) of DEGs versus the distance between their TSS and the nearest loop domain anchor. p = 0.0025, Spearman correlation. (H) Bar graphs of the percentage of genes at binned distances from the TAD boundary that are differentially expressed by >2 log_2_(fold change) upon auxin treatment in the HCT116-RAD21-AID cells. Enrichment of TSSs within 2.5kb or 5kb of boundary not significant by Fisher’s exact test (p = 0.33, p = 0.56, respectively). (I) Bar graphs of the percentage of genes at binned distances from loop anchors that are differentially expressed by >2 log_2_(fold change) upon auxin treatment in the HCT116-RAD21-AID cells. * p = 0.002, ** p < 0.001, Fisher’s exact test. (J) Bar graphs of the percentage of genes at binned distances from loop domain anchors that are differentially expressed by >2 log_2_(fold change) upon auxin treatment in the HCT116-RAD21-AID cells. ** p < 0.001, Fisher’s exact test.

Several differentially expressed genes (DEGs) were in close proximity to TAD boundaries. For example, the transcription start sites (TSSs) of three DEGs, *ID2*, *RNA5SP156*, and *HS3ST1*, are within 16-31 kb of a TAD boundary despite their entire domains being 250 kb-1.98 Mb in size (**Figure 4C**). To test whether this enrichment was true genome-wide and correlated with the change in gene expression, we plotted the distance between each expressed gene (TSS) and the nearest boundary as a function of their log_2_ fold-change in expression. In addition to TAD boundaries, we also assayed anchors of loops and loop domains as defined by Rao et al. (Rao et al., 2017) to examine different types of cohesin-dependent structures separately. Note that these are overlapping definitions and that many TAD boundaries are also loop anchors as is the case for the *RNA5SP156* and *HS3ST1-*proximal boundaries shown in Figure 4C. Overall, TSSs of expressed genes were found at varying distances from architectural boundaries and spanned the full length of the domain size range (**Figures 4E-G**). However, DEGs with a higher fold change in expression were enriched near domain boundaries and loop anchors as compared to lower fold change or unchanged genes, regardless of their structure type and size (p<0.005; **Figures 4E-G**). For example, most DEGs with >2-fold change in expression following cohesin loss were located within 50 kb of a loop anchor, despite an average loop size of ∼350 kb. Importantly, there was no difference in the size of TADs, loops, or loop domains harboring either DEGs or nonDEGs (**Figures S6A-C**). This suggests that the extent of misexpression for DEGs is correlated to their distance from a boundary.

Next, to determine if the most affected DEGs are enriched near TAD boundaries compared to all expressed genes, we calculated the fraction of all expressed genes with >2-fold change in expression at stratified distances from domain boundaries and loop anchors. We found a significant enrichment of DEGs within 2.5 and 5.0 kb of a loop or loop domain anchor (**Figures 4I-J and S6E-F**). In contrast, DEGs showed a bimodal distribution of enrichment in TADs with peaks at 2.5 kb and 100 kb, the latter of which likely reflects inner loop and loop domain anchors (**Figures 4H**). Together, this indicates that genes at architectural boundaries and loop anchors are more likely to be misexpressed to a larger extent following the acute depletion of cohesin in HCT-116 cells.

### Boundary sensitivity is a general signature of cohesin loss in Cornelia de Lange syndrome

To determine if boundary sensitivity is a general signature of cohesin loss, we next turned to patient cells with mutations in NIPBL, which causes a rare developmental disorder called Cornelia de Lange syndrome (CdLS) (Dorsett and Krantz, 2009). Lymphoblastoid cells (LCLs) derived from patients with NIPBL mutations exhibit decreased chromatin-bound cohesin levels and widespread yet modest changes in gene expression (Liu et al., 2009). In particular, 1,501 genes were found to be recurrently differentially expressed across multiple patients (Liu et al., 2009).

Approximately 70% of the 1,501 DEGs were unique to CdLS-derived LCLs and were not differentially expressed in the HCT-116 cell line following auxin treatment, suggesting that many DEGs and associated boundary sites are cell-type-specific (**Figure 5A**). Indeed, only ∼40% of TAD boundary sites were shared between the cell types (Figure 5B and Table S3). We plotted the fold change of CdLS DEGs against the nearest TAD boundary or loop anchor called in the LCL line GM12878 (**Figures 5C-E**) (Rao et al., 2014). Similar to the RAD21 degron system in HCT-116 cells, the extent of misexpression for CdLS-associated DEGs is correlated to their distance from a boundary and/or loop anchor (p<0.001). Together, these data suggest that gene expression at the boundaries of architectural domains is especially sensitive to cohesin dysfunction in patient cells.

**Figure 5:**
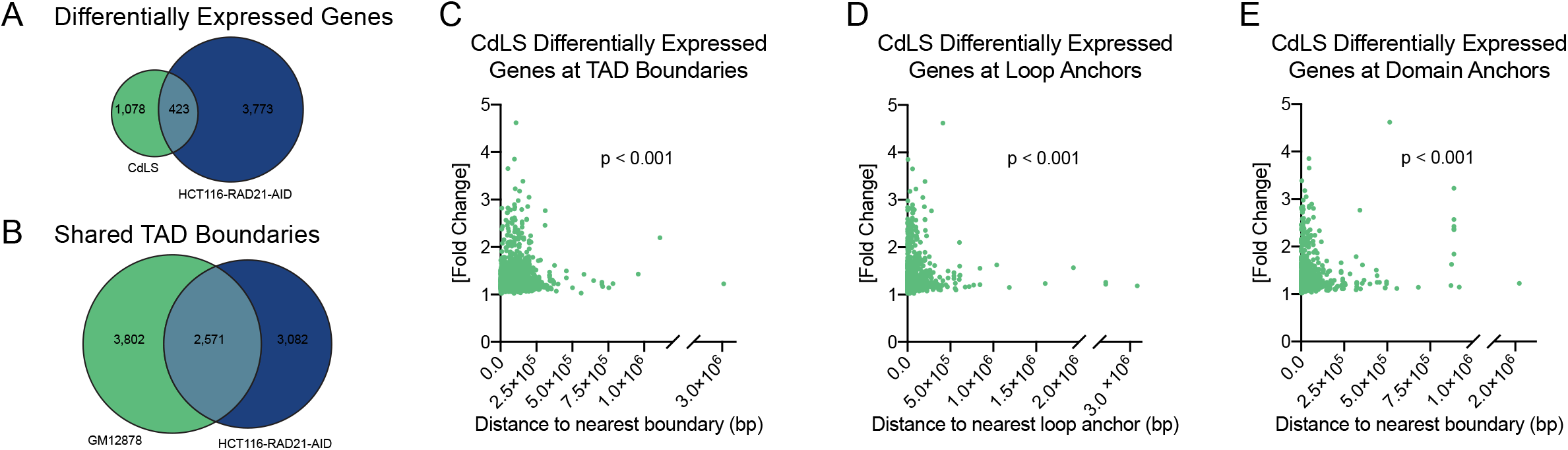
Boundary sensitivity is a general signature of cohesin loss in Cornelia de Lange syndrome. (A) Venn diagram representing overlap between CdLS-associated DEGs and HCT-116-RAD21-AID DEGs. (B) Venn diagram representing the number of TAD boundaries called in GM12878 and HCT-116-RAD21-AID by TopDom, and the number of boundaries shared between the two cell lines. (C) Scatterplot of absolute fold change of CdLS-associated DEGs versus the distance between their TSS and the nearest TAD boundary in GM12878. TADs called by TopDom. p = 0.0002, Fisher’s exact test (enrichment within 2.5kb of TAD boundary). (D) Scatterplot of absolute fold change of CdLS-associated DEGs versus the distance between their TSS and the nearest loop anchor in GM12878. Loop designations from Rao *et al*., (Rao et al., 2014). p = 4.0e-09, Fisher’s exact test (enrichment within 2.5kb of loop anchor). (E) Scatterplot of absolute fold change of CdLS-associated DEGs versus the distance between their TSS and the nearest domain anchor in GM12878. Loop designations from Rao *et al*., (Rao et al., 2014) and called by Arrowhead. p < 2.2e-16, Fisher’s exact test (enrichment within 2.5kb of domain boundary).

### Cohesin alters the topological context and transcriptional bursting frequency of boundary-proximal genes

To further explore the relationship between boundary-proximal DEGs and stochastic domain intermingling, we next tested the arrangement of DEGs relative to their neighboring domains in control and RAD21-depleted cells. We applied three-color FISH to probe the boundary-proximal DEG *CREBL2* along with the upstream portion of its conventional domain and its neighboring downstream domain (**Figure 6A**). We defined four topological configurations based on the position of the gene relative to either domain: 1) the gene locus interacts with its expected contact domain (Domain Maintenance), 2) the gene has switched its contact domain (Domain Switching), 3) the gene no longer interacts with either domain (Domain Exclusion), or 4) the gene interacts simultaneously with both domains (Domain Sharing) (**Figure 6B**). Although all four configurations occurred throughout the cell population, <1% of cells exhibited a domain sharing configuration in which the gene interacted with both domains simultaneously. This indicates that stochastic domain intermingling is not due to complete domain merging but instead arises from the asymmetric incorporation of boundary-proximal chromatin. Indeed, the *CREBL2* gene interacted with the upstream portion of its expected domain in only 21% of chromosomes and interacted exclusively with the neighboring domain at a similar frequency (35%; **Figure 6C**). Most commonly, the gene locus was looped out and excluded from either domain (43%). Similar results were obtained for the boundary-proximal DEG *MCM5* on chromosome 22 (**Figures 6A,D**). These results further highlight the variable nature of domain boundaries and suggest that genes near boundaries are often located outside of their expected topological context.

**Figure 6.**
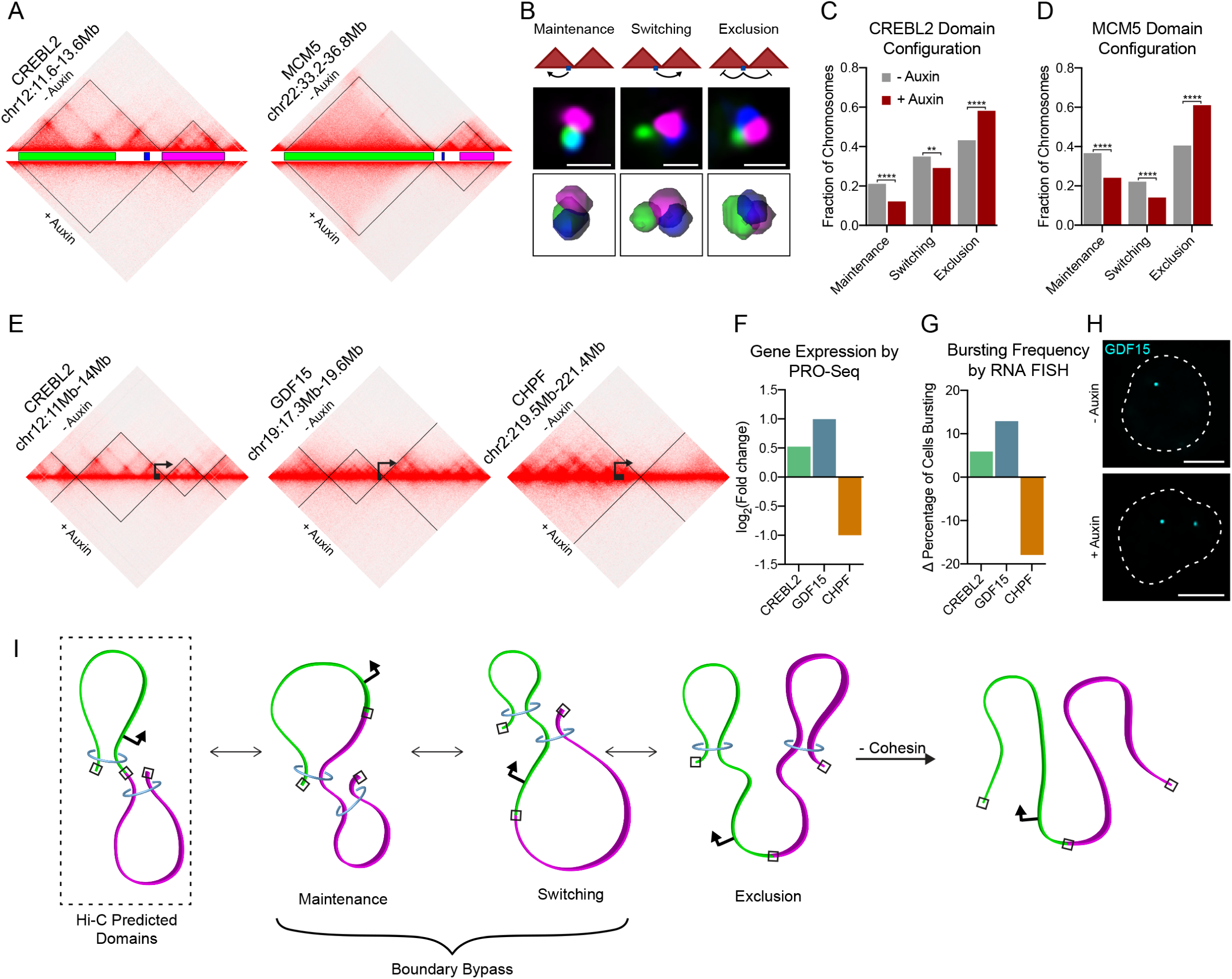
Cohesin alters the topological context and transcriptional bursting frequency of boundary-proximal genes. (A) Hi-C contact matrix of chr12:11.6-13.6Mb (left) and chr22:33.2-36.8Mb (right) and Oligopaint design for each corresponding to (B-D). (B) Cartoon representations for three possible interactions between boundary-proximal genes and their neighboring domains: domain maintenance, switching, and exclusion (top). Representative deconvolved widefield images of three-color FISH to the chr12:11.6-13.6Mb locus illustrating the three domain configurations (middle); scale bar equals 1 μm. Corresponding 3D segmentation of FISH signals below each image. (C) Bar graph representing frequencies of domain configurations at chr12:11.6-13.6Mb. **** p < 0.0001, ** p = 0.0046, Fisher’s exact test. n > 979 chromosomes. (D) Bar graph representing frequencies of domain configurations at chr22:33.2-36.8Mb. **** p < 0.0001, Fisher’s exact test. n > 863 chromosomes. (E) Hi-C contact matrix and relative gene annotation for the *CREBL2*, *GDF15*, and *CHPF* loci. (F) Bar graph depicting gene expression changes by PRO-Seq (Rao et al., 2017). (G) Bar graph depicting bursting frequency changes of each gene following auxin treatment by intronic RNA FISH. n > 327 chromosomes. (H) Representative widefield images for RNA FISH to the *GDF15* gene with one (above) or two (below) bursting alleles. Dashed lines represent nuclear edges. Scale bar equals 5μm. (I) Model depicting loop domain folding patterns inferred by Hi-C and FISH data. Neighboring domains are painted in green and magenta, with Hi-C defined boundaries indicated with a black box, boundary proximal gene promoters represented by arrows, and cohesin shown as a blue ring. Hi-C data predicts neighboring loop domains are spatially separate structures tethered at the boundaries. Imaging shows high levels of inter-TAD interactions, which may occur if cohesin dynamically extrudes through a Hi-C defined boundary and incorporates chromatin from one domain with its neighboring domain. Boundary bypass may occur in either direction (depicted as maintenance and switching). In addition, boundary proximal genes may be looped out from either domain (exclusion) in a subset of cells. In the absence of cohesin, we see an increase in boundary exclusion from its neighboring domains. Depending on the context of the locus, this may lead to either decreased or increased transcription if the boundary-proximal gene were looped away from an active enhancer or repressor, respectively.

Following auxin treatment to deplete RAD21, the *CREBL2* locus was more frequently looped out and excluded from either domain (58% of chromosomes; **Figure 6C**). A similar increase in domain exclusion was found for *MCM5* (**Figure 6D**). At both loci, this was accompanied by a significant reduction in both domain maintenance and switching. Therefore, in the absence of cohesin, domain separation preferentially induces increased exclusion of boundary-proximal genes from their conventional domain in a fraction of cells.

We reasoned that reduced interactions between boundary-proximal genes and their predicted regulatory domain may alter their engagement with enhancers or repressors in a subset of cells. To determine if altered gene activation rates in a subpopulation of cells can account for their moderate changes in expression, we performed intronic RNA FISH to measure transcriptional bursting frequencies at three DEGs, *CREBL2*, *GDF15*, and *CHPF*, which are normally expressed at detectable levels and located in close proximity to a TAD boundary (**Figure 6E**). All three genes exhibited bursty transcription by RNA FISH with 33, 15, and 57% of active alleles, respectively. By PRO-seq analysis, the three genes were previously shown to be up- or downregulated following cohesin degradation (**Figure 1F**) (Rao et al., 2017).

We observed altered bursting frequency of all three genes following RAD21 degradation; a 6% and 13% increase at the *CREBL2* and *GDF15* loci, respectively, and an 18% decrease at *CHPF* (**Figure 6G-H**). The bursting size and intensity were also significantly altered in the two upregulated genes (*CREBL2* and *GDF15*) (**Figures S6G-H**). Remarkably, the shifts in bursting dynamics within the cell population were consistent with their direction and relative changes in expression as shown by PRO-seq (Figure 6F-G). They also mimicked the increased fraction of cells that exhibited domain exclusion of boundary-proximal loci following cohesin loss (**Figures 6C-D**). Together, these results suggest that cohesin promotes proper transcriptional bursting dynamics preferentially at genes near domain boundaries.

## Discussion

Current models suggest that TADs are functional regulons, which spatially separate into 3D structures to insulate gene regulation. However, an emerging theme from recent single-cell based assays is that genome packaging is extremely dynamic and heterogeneous across a cell population (Bintu et al., 2018; Cardozo Gizzi et al., 2019; Finn et al., 2019; Flyamer et al., 2017; Nir et al., 2018). Yet, the frequency, extent, and nature of these heterogeneous structures remains unclear. Using Oligopaint FISH technology to precisely target entire genomic domains defined by Hi-C, we find that, on average, ∼60% of alleles show some degree of intermingling between neighboring TADs. The simplicity of our TAD FISH assay also allowed us to extend this conclusion to multiple loci of different chromatin types, suggesting variable and extensive interactions between neighboring TADs are a widespread feature of the human genome.

The loop extrusion model, in which cohesin complexes extrude DNA until halted by a pair of CTCF motifs, has been proposed to explain the formation of loops and TADs at the population level (Goloborodko et al., 2016; Sanborn et al., 2015). Our results further support this model given the distance-dependent nature and enrichment of inter-TAD interactions near shared boundaries. However, our data further suggest that boundaries are semi-permissible and are bypassed by cohesin in a fraction of cells. Indeed, we find that a fraction of inter-TAD contact frequencies are dependent on cohesin. We also find increased interactions between TADs following WAPL or CTCF knockdown, consistent with WAPL and CTCF restricting cohesin-based chromatin extrusion (Haarhuis et al., 2017; Hansen et al., 2017). Using three-color FISH to examine the configuration of boundary-proximal chromatin, our data further suggests that boundaries are bypassed in an asymmetric fashion, leading to alternating incorporation of boundary-proximal chromatin with neighboring domains. Combined with the dynamic nature of cohesin and CTCF binding (Hansen et al., 2017; Rao et al., 2017), we propose that stochastic boundary bypass toggles boundary incorporation between neighboring domains across a single cell cycle (**Figure 6I**).

It is important to note that cohesin loss does not eliminate inter- or intra-TAD interactions. Despite ∼95% loss of chromatin-bound RAD21, many chromosomes still showed a significant level of inter-TAD and intra-TAD intermingling at all loci tested. In fact, intra-TAD contact frequencies remained higher than inter-TAD contact in the absence of cohesin. This indicates that some aspects of domain and subdomain interactions are not cohesin-dependent. These interactions may simply reflect polymer dynamics and the tendency of neighboring regions to spontaneously cluster together (Fudenberg et al., 2016; Mirny et al., 2019). Alternatively, the variegated spreading of histone modifications could lead to variable interactions across boundaries and self-association via phase separation (Elgin and Reuter, 2013; Hnisz et al., 2017; Sabari et al., 2018; Strom et al., 2017). Regardless, this result is consistent with the identification of ‘TAD-like’ contact domains across a single locus in the absence of cohesin as revealed by sequential FISH (Bintu et al., 2018) and further suggests that this is a widespread phenomenon. The nature of these interactions and their functional relevance, if any, will be an interesting direction for future investigation.

Finally, we consider our results in the context of gene expression and the role TADs are proposed to play in gene regulation. Many studies have suggested that loop domains formed between CTCF and cohesin binding sites create insulated regulatory neighborhoods, partially protecting genes from the influence of enhancers outside their typical domain (Flavahan et al., 2016; Lupianez et al., 2015). Our study shows that inter-TAD contacts are most likely not a deterministic factor in ectopic gene activation or repression given the frequency and extent of interactions between neighboring domains. Instead, CTCF and boundaries, in general, may insulate gene regulation not strictly through spatial separation, but possibly through a change in the type, stability, or combination of interactions within a domain. What then is the role of cohesin-mediated inter-TAD interactions in gene regulation? We discovered a novel signature of cohesin loss in which genes near boundary elements are more likely to be misexpressed to a larger extent. We find a similar signature in cells derived from NIPBL-deficient individuals, suggesting that this is a feature of classical Cornelia de Lange Syndrome. Combined with our findings that boundary-proximal chromatin is more often excluded from neighboring domains following cohesin loss, we propose that cohesin promotes the alternating incorporation of boundary-proximal chromatin with either of its neighboring domains to ensure a high probability of contact between all portions of a TAD (**Figure 6I**). Thus, population-based reproducibility in gene expression may be achieved through stochastic interactions across domain boundaries. This would be especially important for boundary-proximal genes, as these genes would need to travel up to twice the maximum distance a gene near the TAD center would to contact a distal regulatory element. Indeed, increased exclusion of these genes from neighboring domains following cohesin loss could explain both downregulation and upregulation of DEGs if a boundary-proximal gene were looped out away from a distal enhancer or repressor, respectively. Thus, our results provide one explanation for heterogeneous TAD organization and suggest a new paradigm in which cohesin dynamically bypasses boundaries to create stochastic domain structures that help regulate gene expression.

## Methods

### TopDom TAD calling

Published Hi-C data was downloaded from GEO database (GSE104334, GSE63525) and 25kb-window contact matrix was extracted with the Juicebox tools (v1.9.8, “dump observed KR”). TopDom (v0.0.2) was then used to identify topological domains from the contact matrix, with the optimization parameter window.size set to be 10 (including data within 10 windows when computing local topological domains). TADs were further processed to include only those between 250kb and 2Mb in size.

### Compartment analysis

To designate compartments in the HCT-116-RAD21-AID cell line, we determined eigenvectors from Hi-C data published in Rao *et al*. (Rao et al., 2017). We applied the eigenvector feature annotation package included in the Juicer software (https://github.com/aidenlab/juicer/wiki/Eigenvector) with KR normalization at 500 Kb resolution (Durand et al., 2016b). While the sign of the eigenvector typically reflects the A or B compartment designation, we further confirmed compartment designations by comparing the eigenvector coordinates with ChIP-seq data for various histone modifications marking active and inactive chromatin published in (Rao et al., 2017) (Figure S1).

### Oligopaint design and synthesis

To label TADs in the HCT-116-RAD21-AID cell line, we first applied the TopDom TAD algorithm to Hi-C data published in Rao *et al*. (Rao et al., 2017). This method defines TAD boundaries as local minima of contact frequency between up and downstream regions per binned chromatin segment. We then identified RAD21 and CTCF colocalized sites at the boundaries between TADs from ChIP-seq data available on ENCODE (ENCSR000BSB and ENCSR000DTO, respectively). We defined the TAD boundary coordinates as the center of the corresponding RAD21 ChIP-seq peak.

We then applied the OligoMiner design pipeline to design DNA FISH probes to the coordinates found in Table S2 (Beliveau et al., 2018). Oligopaints were designed to have 42 bases of homology with an average of five probes per kb and were purchased from CustomArray. Additional bridge probes were designed to the sub-TADs and boundary-proximal genes to amplify the signal (Beliveau et al., 2015). Oligopaints were synthesized as previously described (Rosin et al., 2018).

### Cell culture

HCT-116-RAD21-AID cells were obtained from Natsume et al. (Natsume et al., 2016). The cells were cultured in McCoy’s 5A media supplemented with 10% FBS, 2 mM L-glutamine, 100 U/ml penicillin, and 100ug/ml streptomycin at 37°C with 5% CO2. Cells were selected with 100 ug/ml G418 and 100 ug/ml HygroGold prior to experiments.

To avoid heterogeneity due to cell cycle, the HCT-116-RAD21-AID cells were synchronized at the G1/S transition. First, to arrest cells in the S-phase, cells were grown in media supplemented with 2mM thymidine (Sigma-Aldrich T1895) for 12 hours. Cells were then resuspended in media and allowed to grow for 12 hours to exit S-phase. To arrest at the G1/S transition, we grew cells in media supplemented 400 uM mimosine (Sigma-Aldrich M025) for 12 hours. Lastly, we replaced media with either 400 uM mimosine + 500 uM indole-3-acetic acid (auxin; Sigma-Aldrich I5148) supplemented media to degrade RAD21 or 400 uM mimosine-supplemented media alone as an untreated control; cells were incubated with or without auxin for 6 hours then harvested for experiments. Synchronization was confirmed by IF (Figure S2).

### RNAi

HCT-116 cells were cultured in McCoy’s 5A media supplemented with 10% FBS, 2 mM L-glutamine, 100 U/ml penicillin, and 100ug/ml streptomycin at 37°C with 5%CO2. The following siRNAs (Dharmacon) were used: Non-targeting control (D-001210-05-05), NIPBL (J-012980-08; target sequence: 5’-CAACAGAUCACAUAGAGUU-3’), WAPL (L-026287-01-0005; target sequences administered as a pool: 5’-GGAGUAUAGUGCUCGGAAU-3’, 5’-GAGAGAUGUUUACGAGUUU-3’, 5’-CAAACAGUGAAUCGAGUAA-3’, 5’-CCAAAGAUACACGGGAUUA-3’), CTCF (L-020165-00-0010; target sequences administered as a pool: 5’-GAUGAAGACUGAAGUAAUG-3’, 5’-GGAGAAACGAAGAAGAGUA-3’, 5’-GAAGAUGCCUGCCACUUAC-3’, 5’-GAACAGCCCAUAAACAUAG-3’). Duplex siRNA were incubated for 20 minutes at room temperature with RNAiMAX transfection reagent (ThermoFisher) in Opti-MEM reduced serum media (ThermoFisher) and seeded into wells. HCT-116 were trypsinized and resuspended in antibiotic free media, then plated onto siRNA for a final siRNA concentration of 50 nM. After 72h (NIPBL, WAPL, non-targeting control) or 96h of RNAi treatment (CTCF), cells were harvested for experiments.

### DNA Fluorescence in situ hybridization (FISH)

Cells were settled on poly-L-lysine treated glass slides for two hours, or uncoated high precision 22×22 mm coverslips for 6 hours. Cells were then fixed to the slide or coverslip for 10 minutes with 4% formaldehyde in PBS with 0.1% Tween 20, followed by three washes in PBS for 5 minutes each. Slides and coverslips were stored in PBS at 4°C until use.

For experiments imaged by widefield microscopy, FISH was performed on slides. Slides were warmed to room temperature (RT) in PBS for 10 minutes. Cells were permeabilized in 0.5% Triton-PBS for 15 minutes. Cells were then dehydrated in an ethanol row, consisting of 2 minute incubations in 70%, 90%, and 100% ethanol consecutively. The slides were then allowed to dry for 2 minutes at room temperature. Slides were incubated 5 minutes each in 2xSSCT (0.3 M NaCl, 0.03 M sodium citrate, 0.1% Tween-20) and 2xSSCT/50% formamide at room temperature, followed by a 1 hour incubation in 2xSSCT/50% formamide at 37°C. Hybridization buffer containing primary Oligopaint probes, hybridization buffer (10% dextran sulfate/2xSSCT/50% formamide/4% polyvinylsulfonic acid (PVSA)), 5.6 mM dNTPs, and 10 ug RNaseA was added to slides, covered with coverslip, and sealed with rubber cement. 50 pmol of probe was used per 25 ul hybridization buffer. Slides were then denatured on a heat block in a water bath set to 80°C for 30 minutes, then transferred to a humidified chamber and incubated overnight at 37°C. The following day, coverslips were removed and slides were washed in 2xSSCT at 60°C for 15 minutes, 2xSSCT at RT for 10 minutes, and 0.2xSSC at RT for 10 minutes. Next, hybridization buffer (10% dextran sulfate/2xSSCT/10% formamide/4% polyvinylsulfonic acid (PVSA)) containing secondary probes conjugated to fluorophores (10 pmol/25 ul buffer) was added to slides, covered with coverslip, and sealed with rubber cement. Slides were placed in humidified chamber and incubated 2 hours at RT. Slides were washed in 2xSSCT at 60°C for 15 minutes, 2xSSCT at RT for 10 minutes, and 0.2xSSC at RT for 10 minutes. To stain DNA, slides were washed with Hoechst (1:10,000 in 2xSSC) for 5 minutes. Slides were then mounted in SlowFade Gold Antifade (Invitrogen). For experiments imaged by 3D-STORM, FISH was performed on coverslips as described above for slides, without DNA staining.

### Immunofluorescence

Slides were prepared as for DNA FISH. Cells were permeabilized in 0.1% Triton-PBS for 15 minutes, then washed three times in PBS-T (PBS with 0.1% Tween 20) for 10 minutes each. Proteins were blocked in 1% bovine serum albumin (BSA) in PBS-T for 1 hour at RT. Primary antibodies diluted in 1% BSA-PBS-T were added to the slide, covered with coverslip, and sealed with rubber cement. Slides were transferred to humidified chamber and incubated overnight at 4°C. The following day, slides were washed three times in PBS-T for 10 minutes each. Secondary antibody was diluted in 1% BSA-PBS-T, added to slide, covered with coverslip, and sealed with rubber cement. Slides were transferred to humidified chamber and incubated at RT for 2 hours. Slides were then washed twice in PBS-T for 10 minutes each, and once in PBS for 10 minutes. To stain DNA, slides were washed with Hoechst (1:10,000 in 2xSSC) for 5 minutes. Slides were then mounted in SlowFade Gold Antifade (Invitrogen).

### Widefield microscopy, image processing, and data analysis

Images were acquired on a Leica wide-field fluorescence microscope, using a 1.4 NA 63x oil-immersion objective (Leica) and Andor iXon Ultra emCCD camera. All images were deconvolved with Huygens Essential version 18.10 (Scientific Volume Imaging, The Netherlands, http://svi.nl), using the CMLE algorithm, with SNR:20-40 and 40 iterations. The deconvoluted images were segmented and measured using a modified version of the TANGO 3D-segmentation plug-in for ImageJ (Ollion et al., 2013, 2015; Rosin et al., 2018). Edges of nuclei and FISH signals were segmented using a Hysteresis-based algorithm. Contact between signals was defined by either ≥ 2.5% overlap or two objects with greater than ≥ 500 nm^3^ voxel colocalization.

### 3D-STORM Imaging

3D-STORM images were acquired on a Bruker Vutara 352 super-resolution microscope with an Olympus 60x/1.2 NA W objective and Hamamatsu ORCA Flash 4.0 v3 sCMOS camera. The 640 and 561 lasers were used to acquire images for TADs labeled with Alexa 647 and CF568 respectively. 3D-STORM imaging buffer contained 10% glucose, 2xSSC, 0.05M Tris, 2% glucose oxidase solution, and 1% betemercaptoethanol. The glucose oxidase solution consisted of 20 mg/mL glucose oxidase and 2 mg/mL catalase from bovine liver dissolved in buffer (50 mM Tris and 10 mM NaCl).

Fields of view were selected by widefield such that each nucleus contained two distinct pairs of TADs. Z-stacks were determined such that both homologs were within the imaged space, and ranged from 3.6-9.6 um. Localizations were then recorded in 0.1 um steps; 150 frames were recorded per z step, and the z stack was cycled through 3-4 times. Imaging between channels was carried out sequentially, with the Alexa647 probe image first. Imaging of the CF568 fluorophore was supplemented with 0.5% power of the 405 laser at the second to last cycle. Localization of the fluorophores was carried out using the B-SPLINE PSF interpolation spline.

Images were further filtered for localizations with <20 nm and <30 nm radial precision for Alexa647 and CF568 respectively, and <60 nm and <80 nm axial precision for Alexa647 and CF568 respectively. To define the largest clusters of signal, we applied the DBScan algorithm with 0.250 um maximum particle distance, 45 minimum particles to form cluster, and 0.250 um hull alpha shape radius.

### RNA FISH

Custom Stellaris® FISH Probes were designed by utilizing the Stellaris® RNA FISH Probe Designer version 4.2 (Biosearch Technologies, Inc., Petaluma, CA). Intronic probes to CREBL2, CHPF, and GDF15 were synthesized with Quasar 670. Cells were seeded and fixed onto Lab-Tek II 8-well chambered coverglass dishes (Thermo Fisher Scientific), using the same fixation procedures as DNA FISH. After fixation, cells were permeabilized overnight in 300 ul of 70% ethanol containing 2% SDS. The following day, cells were washed in 2x SSC containing 10% formamide for 5 min, and then probes were hybridized with cells in a 200 ul mixture containing 10% dextran sulfate, 2x SSCT, 10% formamide, 4% polyvinylsulfonic acid (PVSA), 2% SDS, 2.8 mM dNTPs, and 15.6 nM of probe. 300 ul of mineral oil was added to each well to prevent evaporation, and the dishes were placed in a humidified chamber at 37°C overnight. The next day, cells were washed in 2x SSC containing 10% formamide twice for 30 min each, with the last wash containing 0.1 ug/ml of Hoechst 33342 stain. Cells were then washed with 2x SSC (no formamide) for 5 min before mounting in 300 ul of buffer containing glucose oxidase (37 ug/ml), catalase (100 ug/ml), 2x SSC, 0.4% glucose, and 10 mM Tris-HCl prior to imaging.

### Subcellular Protein Fractionation and Western Blots

Cells were trypsinized and resuspended in fresh media, washed once in cold DPBS, and then spun for 1200xg for 5 minutes at 4°C. The cell pellet was then either processed with the Subcellular Protein Fractionation Kit for Cultured Cells kit (Thermo Scientific, 78840), or resuspended in 1 x RIPA buffer with protease inhibitors to extract whole cell lysate. The sample was nutated for 30 minutes at 4C, and then spun for 16000xg for 20 minutes at 4°C. Supernatant containing protein was extracted and stored at −80°C.

For western blots, protein was mixed with NuPAGE LDS sample buffer and sample reducing agent (Thermo Fisher Scientific), denatured at 70°C for 10 minutes, then cooled on ice. Benzonase was added to the sample (final concentration 8.3 U/ul), followed by a 15 minute incubation at 37°C. 30 ul of each sample were run on Mini-PROTEAN TGX Stain-Free Precast Gels (Bio-Rad) for about 25 minutes at 35mA per gel. Gel was then activated on ChemiDoc MP Imaging System (Bio-Rad) for 5 minutes. Protein was then transferred to 0.2 um nitrocellulose filter at 100V for 45 minutes. Nitrocellulose filter was then washed twice in TBS (150 nM NaCl, 20 mM Tris) for 5 minutes, and blocked in 5% milk in TBS-T (TBS with 0.05% Tween 20) for 30 minutes. Nitrocellulose filter was washed again twice in TBS-T, then incubated with primary antibody diluted in 5% milk in TBS-T overnight at 4°C. The following day, the nitrocellulose filter was washed twice in TBS-T for 5 minutes each, then incubated with secondary antibodies diluted in 5% milk in TBS-T for 1 hour at room temperature. The nitrocellulose filter was then washed twice in TBS-T for 15 minutes each, followed by a final 15 minute wash in TBS. For blots probed with secondary antibodies conjugated to HRP, stain-free image was acquired then blot was incubated in 1:1 mixture of Clarity Western ECL Substrate reagents (Bio-Rad). Blots were then imaged on ChemiDoc MP Imaging System and analyzed with Bio-Rad Image Lab software (v5.2.1).

### Antibodies

Immunofluorescence was performed using the following primary antibodies: RAD21 (Santa Cruz sc-166973, 1:100), PCNA (Santa Cruz sc-56, 1:100), CENPF (Novus Biologicals NB500-101, 1:100). Secondary antibodies used: Goat anti-Rabbit (Jackson ImmunoResearch 111-165-003, 1:200), Sheep anti-Mouse (Jackson ImmunoResearch 505-605-003, 1:100). Western blots were performed with the following primary antibodies: RAD21 (Abcam ab992, 1:1000), NIPBL (Santa Cruz sc-374625, 1:400), WAPL (Santa Cruz sc-365189, 1:250), Histone H3 (Abcam ab1791, 1:40000), and CTCF (Santa Cruz sc-271474, 1:500). Secondary antibodies used: Goat anti-Rabbit (Jackson ImmunoResearch 111-165-003, 13:10000), Goat anti-Mouse (Jackson ImmunoResearch 115-545-003, 13:10000), Anti-mouse IgG HRP-linked Antibody (Cell Signaling Technologies #7076, 1:5000), Anti-rabbit IgG HRP-linked Antibody (Cell Signaling Technologies #7074, 1:5000).

### Reproducibility and statistical analysis

Each experiment performed in this study was repeated with at least two complete biological replicates and, in some cases, additional technical replicates. The total number of samples (*n*) noted in each experiment is the sum of a single replicate. When necessary, statistical analyses used for each experiment are noted in the figure legends. Statistical tests were performed using either R or Prism 7 software by GraphPad. Figures were assembled in Adobe Illustrator.

## Acknowledgements

We would like to thank Leah Rosin, Melike Lakadamyali, and members of the Joyce lab for helpful discussions and critical reading of the manuscript. Additionally, we thank Masato Kanemaki for the HCT-116-RAD21-AID cell line and Suhas Rao and Erez Lieberman-Aiden for sharing critical primary datasets. We also thank Andrea Stout at the Penn Cell and Developmental Biology Microscopy core and Carl Eberling from Bruker for assistance with super-resolution imaging and analysis. This work was supported by a Charles E Kaufman grant from The Pittsburgh Foundation (KA2017-91787) to E.F.J and NIH grants R35GM128903 to E.F.J. and T32GM008216 to J.M.L.

## Declaration of interests

The authors declare no competing interests.

---

**Figure S1.**
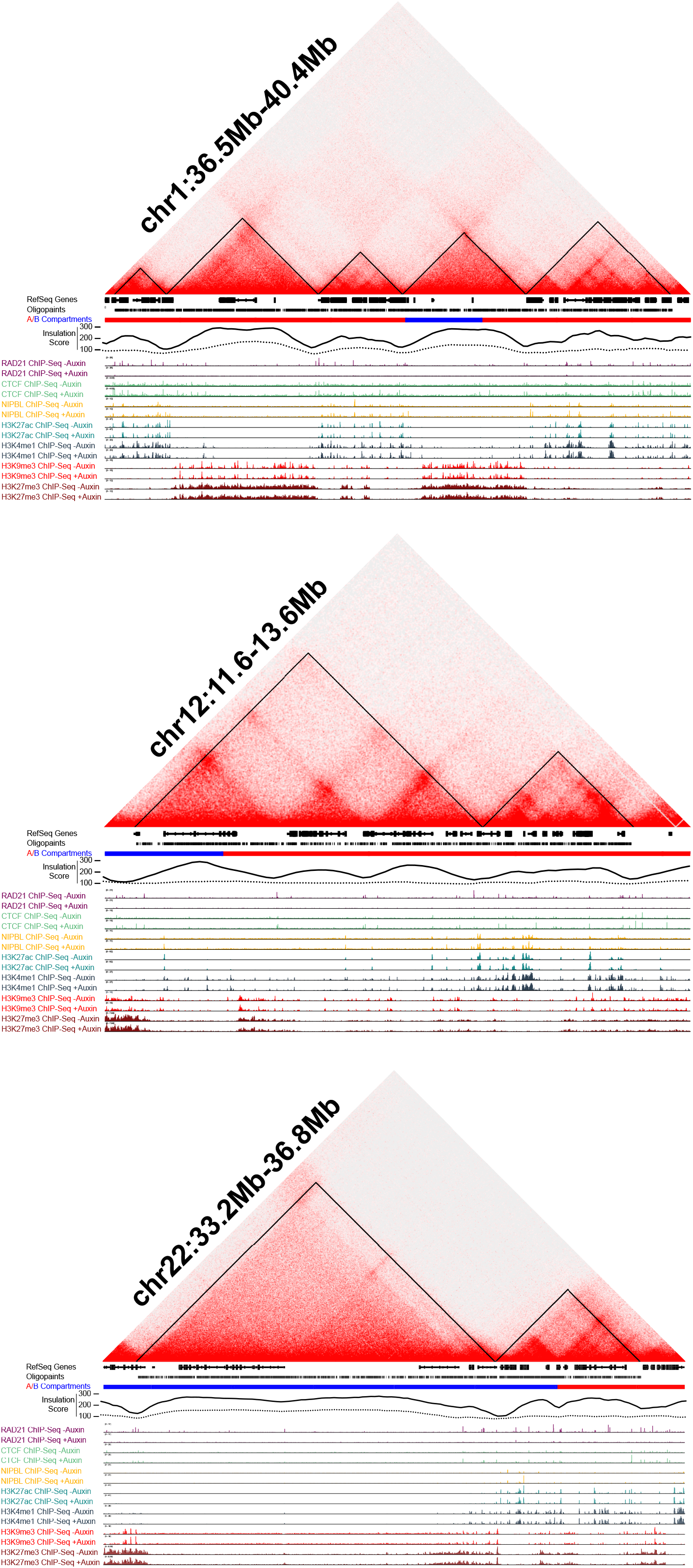
Genomic landscape at Oligopaint target regions. Genomic profiles of loci probed by Oligopaint FISH probes. (Upper panel) chr1:36500000-40400000. (Middle panel) chr12:11600000-13600000. (Lower panel) chr22:33200000-36800000. Hi-C contact matrices for HCT-116-RAD21-AID cell line from (Rao et al., 2017), visualized with Juicebox v1.9.0 (Durand et al., 2016a). Corresponding gene density, compartment designations, and insulation score computed by the TopDom domain algorithm based on mean contact frequency at 25Kb bins. The solid line represents the insulation score computed from the untreated HCT-116-RAD21-AID cell line and the dotted line represents the insulation score computed from the auxin treated HCT-116-RAD21-AID cell line. ChIP-seq tracks published in (Rao et al., 2017) depict protein binding and histone modifications in the HCT-116-RAD21-AID cell line prior to and following 6 hours of auxin treatment to deplete RAD21. Genomic tracks visualized using Integrative Genomics Viewer (Robinson et al., 2011).

**Figure S2.**
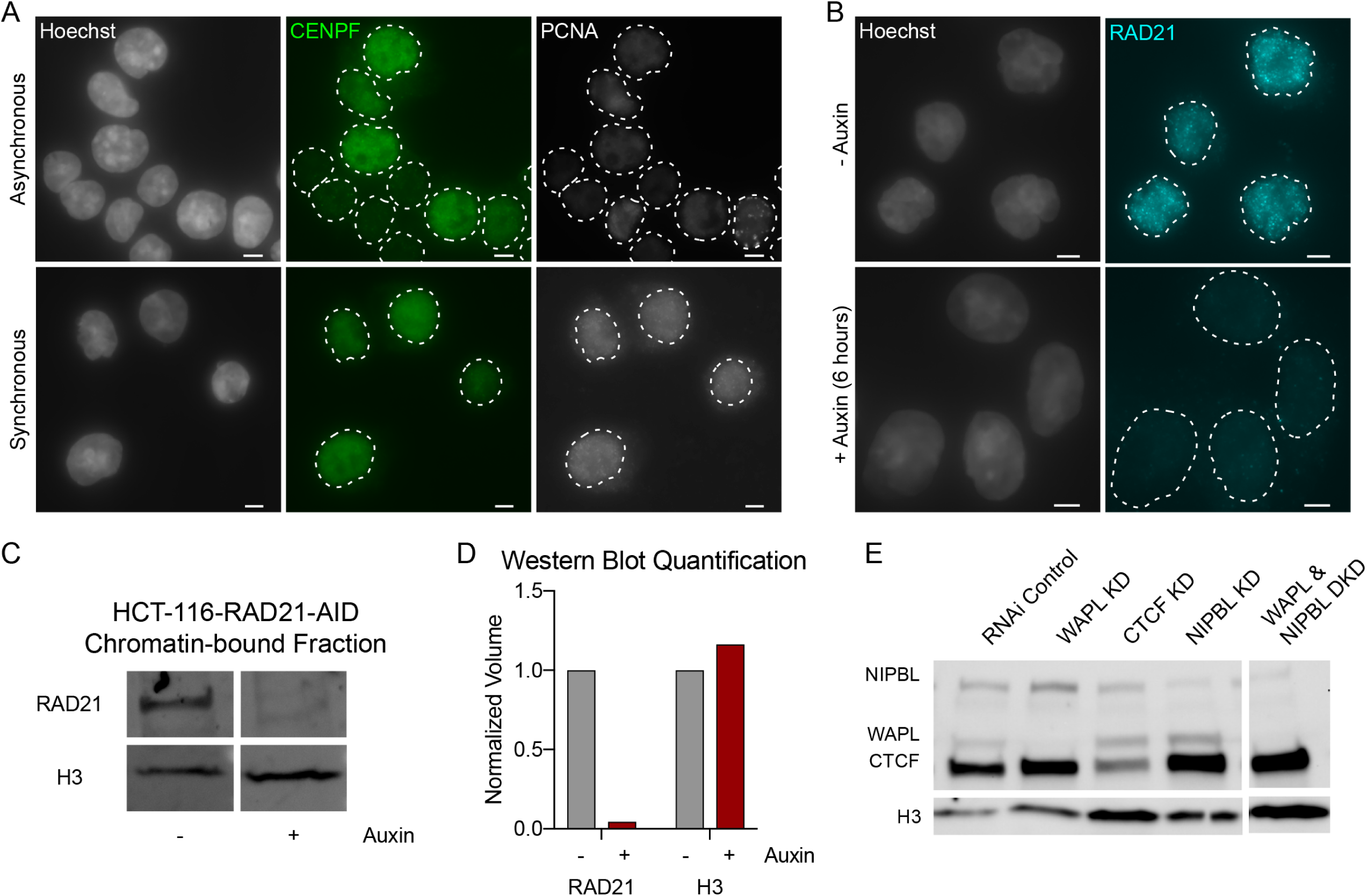
Cell cycle synchronization and knockdowns. (A) HCT-116-RAD21-AID cells were synchronized at the G1/S transition. Immunofluorescence for CENPF (green) to indicate cells in G2 and PCNA (grey) to mark cells in S phase. DNA (Hoescht stain) is shown in grey in first column. Dashed lines represent nuclear edges. Scale bar equals 5 μm. (B) Immunofluorescence for RAD21 (cyan). DNA (Hoescht stain) is shown in grey. Dashed lines represent nuclear edges. Scale bar equals 5 μm. (C) Western blot to RAD21 protein in the chromatin-bound fraction of HCT-116-RAD21-AID cells with no auxin treatment (-) or following 6 hours of auxin treatment (+). Histone H3 as loading control. Protein was labeled using fluorescent secondary antibodies. (D) Fluorescence quantification of RAD21 and H3 isolated from the chromatin-bound fraction of protein corresponding to blot in (C) using Image Lab v5.2.1. Protein intensity normalized to total protein per well (via stain-free technology) and presented as fraction of protein observed in untreated (-auxin) conditions; we observe a 96% reduction in chromatin-bound RAD21 following auxin treatment. (E) Western blot to NIPBL, WAPL, and CTCF protein in whole cell lysate of HCT-116 cells following RNAi. Histone H3 as loading control. Protein was labeled using HRP-conjugated secondary antibodies.

**Figure S3.**
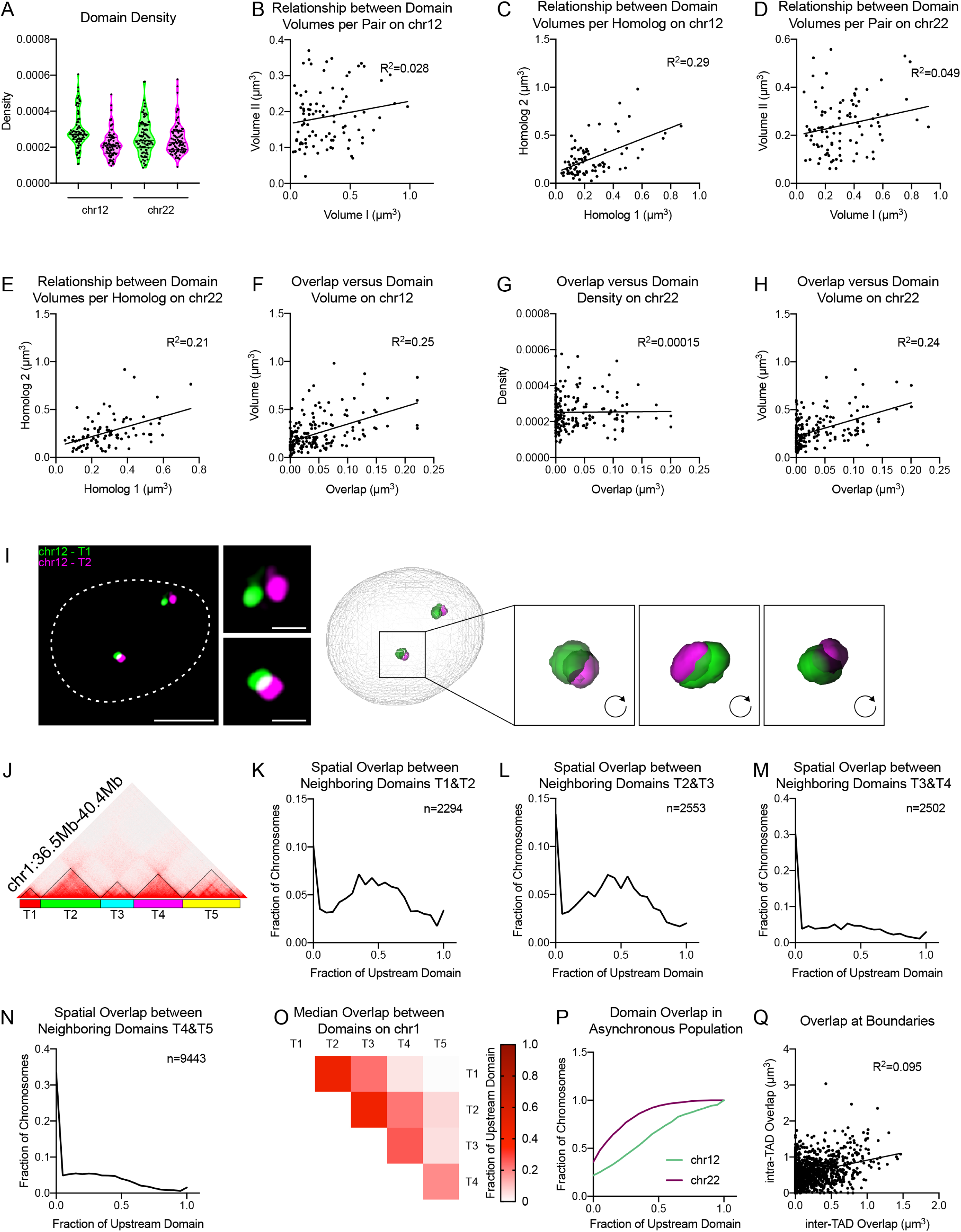
Additional information related to Figure 1. (A) Violin plots of density of localizations per volume per domain as quantified from 3D-STORM images. n > 90 chromosomes. (B) Scatterplot depicting the relationship between domain volumes on the same chromosome by 3D-STORM on chr12:11.Mb-13.6Mb. n = 91 chromosomes. (C) Scatterplot depicting the relationship between domain volumes between homologs by 3D-STORM on chr12:11.Mb-13.6Mb. n = 82 chromosomes. (D) Scatterplot depicting the relationship between domain volumes on the same chromosome by 3D-STORM on chr22:33.2Mb-36.8Mb. n = 95 chromosomes. (E) Scatterplot depicting the relationship between domain volumes between homologs by 3D-STORM on chr22:33.2Mb-36.8Mb. n = 86 chromosomes. (F) Scatterplot of overlap volume (x-axis) versus domain volume (y-axis) by 3D-STORM for the chr12:11.6Mb-13.6Mb locus. n = 91 chromosomes. (G) Scatterplot of overlap volume (x-axis) versus domain density (y-axis) by 3D-STORM for the chr22:33.2Mb-36.8Mb locus. n = 95 chromosomes (H) Scatterplot of overlap volume (x-axis) versus domain volume (y-axis) by 3D-STORM for the chr22:33.2Mb-36.8Mb locus. n = 95 chromosomes. (I) Representative deconvolved widefield image of FISH to neighboring TADs on chr12 with corresponding 3D reconstruction by TANGO image analysis software. (J) Hi-C contact matrix and Oligopaint design corresponding to (K-O). (K) Histograms of overlap between the neighboring domains T1 & T2. Overlap normalized to the volume of the upstream domain. n = 2294 chromosomes. (L) Histograms of overlap between the neighboring domains T2 & T3. Overlap normalized to the volume of the upstream domain. n = 2553 chromosomes. (M) Histograms of overlap between the neighboring domains T3 & T4. Overlap normalized to the volume of the upstream domain. n = 2502 chromosomes. (N) Histograms of overlap between the neighboring domains T4 & T5. Overlap normalized to the volume of the upstream domain. n = 9443 chromosomes. (O) Heatmap indicating median spatial overlap between domains on chr1:35.6Mb-40.4Mb. Overlap normalized to the volume of the upstream domain. n > 2294 chromosomes. (P) Cumulative frequency distribution of spatial overlap between neighboring domains on chr12:11.6Mb-13.6Mb and chr22:33.2Mb-36.8Mb in asynchronous HCT-116 cells. Overlap normalized to the volume of the upstream domain. n > 716 chromosomes. (Q) Scatterplot of inter-TAD overlap versus intra-TAD overlap at the chr22:33.2-36.8Mb locus. Overlap depicted as fraction of the boundary-proximal subTAD volume. n = 1060 chromosomes. See Figure 1J for corresponding Oligopaint design.

**Figure S4.**
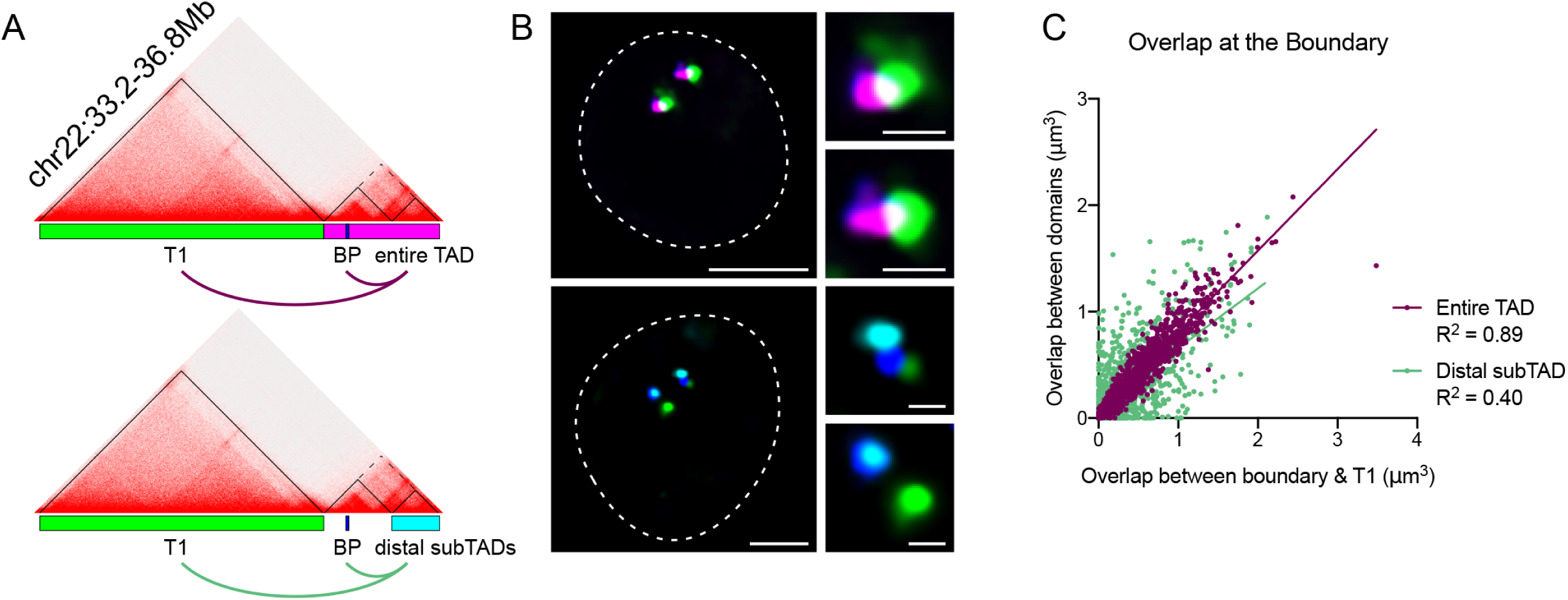
Additional information related to Figure 2. (A) Hi-C contact matrix and Oligopaint designs corresponding to (B-C). (B) Representative deconvolved widefield images. Dashed lines represent nuclear edges. Scale bar equals 5 μm (left) or 1 μm (zoomed images, right). (C) Scatterplot of spatial overlap between the boundary-proximal probe and upstream domain (x-axis) and the entire downstream TAD (purple) or distal subTAD (green). n > 922 chromosomes.

**Figure S5.**
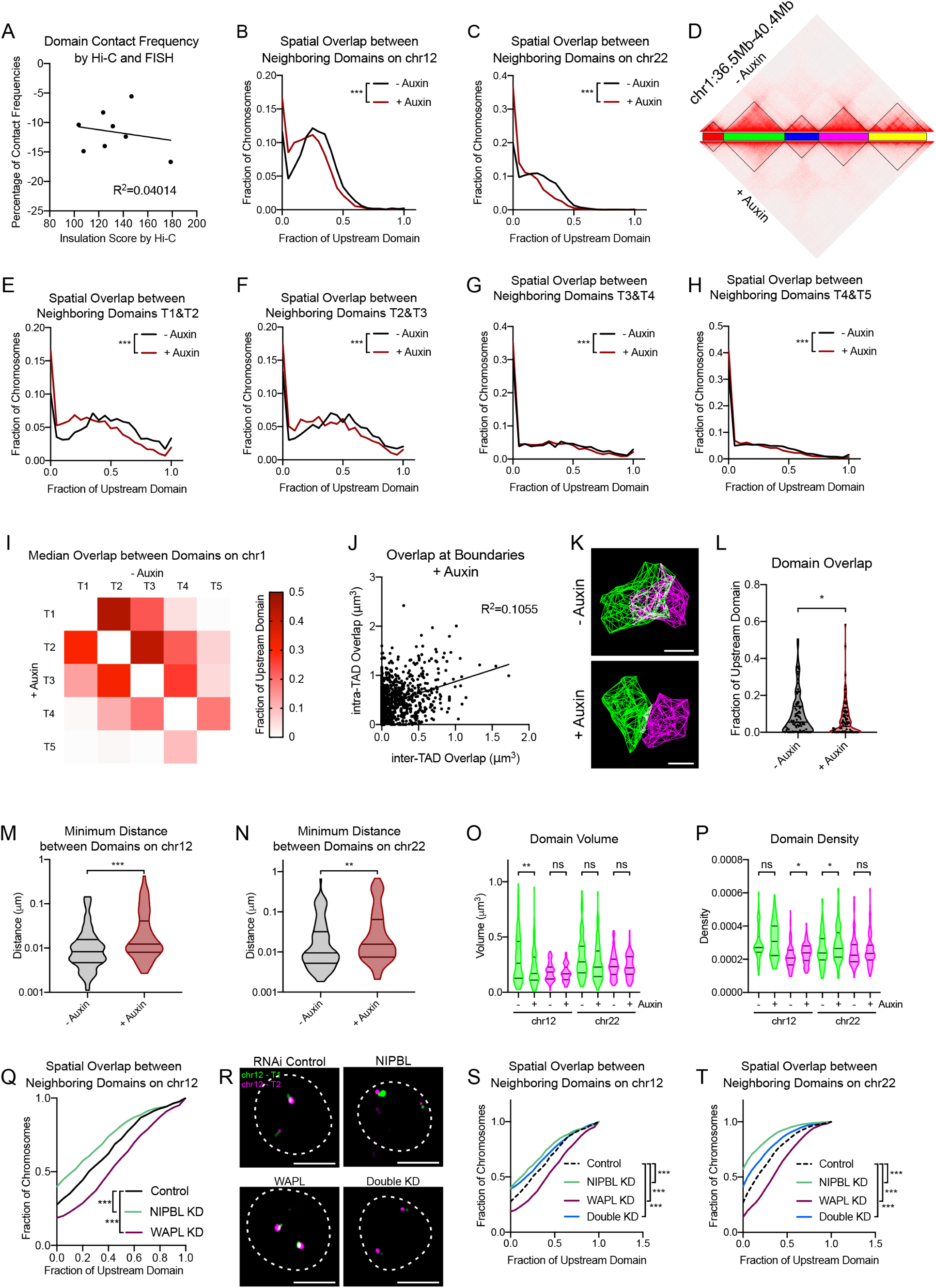
Additional information related to Figure 3. (A) Scatterplot of boundary insulation score determined by TopDom analysis of Hi-C versus the difference in percent contact frequencies between domains following auxin treatment by FISH. Each point represents the average of two biological replicates for eight neighboring domain pairs. (B) Histogram of spatial overlap between domains on chr12:11.6Mb-13.6Mb, normalized to the volume of the upstream domain. p < 0.001, two-tailed Mann Whitney test. n > 3661 chromosomes. (C) Histogram of spatial overlap between domains on chr22:33.2Mb-36.8Mb, normalized to the volume of the upstream domain. p < 0.001, two-tailed Mann Whitney test. n > 2803 chromosomes. (D) Hi-C contact matrix and Oligopaint design corresponding to (E-I). (E) Histograms of overlap between the neighboring domains T1 & T2, normalized to the volume of the upstream TAD. *** p < 0.001, two-tailed Mann Whitney test. n > 2175 chromosomes. (F) Histograms of overlap between the neighboring domains T2 & T3, normalized to the volume of the upstream TAD. *** p < 0.001, two-tailed Mann Whitney test. n > 2353 chromosomes. (G) Histograms of overlap between the neighboring domains T3 & T4 normalized to the volume of the upstream TAD. *** p < 0.001, two-tailed Mann Whitney test. n > 2488 chromosomes. (H) Histograms of overlap between the neighboring domains T4 & T5, normalized to the volume of the upstream TAD. *** p < 0.001, two-tailed Mann Whitney test. n > 8695 chromosomes. (I) Heatmap indicating median spatial overlap between domains on chr1:35.6Mb-40.4Mb. Overlap normalized to the volume of the upstream TAD. n > 2140 chromosomes. (J) Scatterplot of inter-TAD overlap versus intra-TAD overlap at the chr22:33.2-36.8Mb locus following auxin treatment. Overlap depicted as fraction of the boundary-proximal subTAD volume. n > 1060 chromosomes. See Figure 1J for corresponding Oligopaint design. (K) Representative 3D-STORM images of neighboring TADs on chr22:33.2Mb-36.8Mb. Scale bar equals 500 nm. (L) Violin plots of spatial overlap between domains, normalized to volume of the upstream domain. * p = 0.029, two-tailed Mann Whitney test. n > 95 chromosomes. (M) Violin plot of minimum distance between localizations contained within each domain on chr12:11.6Mb-13.6Mb as measured from 3D-STORM data. *** p < 0.001, two-tailed Mann Whitney test. n > 76 chromosomes. (N) Violin plot of minimum distance between localizations contained within each domain on chr22:33.2Mb-36.8Mb as measured from 3D-STORM data. ** p = 0.009, two-tailed Mann Whitney test. n > 95 chromosomes. (O) Violin plots of domain volumes as measured from 3D-STORM data. ** p = 0.008, two-tailed Mann Whitney test. n > 76 chromosomes. (P) Violin plots of domain densities as measured from 3D-STORM data. * p = 0.014 chr12-II, * p = 0.025 chr22-I, two-tailed Mann Whitney test. n > 76 chromosomes. (Q) Cumulative frequency distribution of spatial overlap between neighboring domains on chr12:11.6Mb-13.6Mb in control, NIPBL, and WAPL depleted cells. Overlap normalized to the volume of the upstream domain. *** p < 0.001, two-tailed Mann Whitney test. n > 636 chromosomes. (R) Representative deconvolved widefield images for HCT-116 cells depleted for NIPBL, WAPL, or both by siRNA. Dashed lines represent nuclear edges. Scale bar equals 5 μm. (S) Cumulative frequency distribution of spatial overlap between neighboring domains on chr12:11.6Mb-13.6Mb in control, NIPBL, WAPL, and both NIPBL and WAPL depleted cells. Overlap normalized to the volume of the upstream domain. *** p < 0.001, two-tailed Mann Whitney test. n > 636 chromosomes. (T) Cumulative frequency distribution of spatial overlap between neighboring domains on chr22:33.2Mb-36.8Mb in control, NIPBL, and WAPL depleted cells. Overlap normalized to the volume of the upstream domain. *** p < 0.001, two-tailed Mann Whitney test. n > 1440 chromosomes.

**Figure S6.**
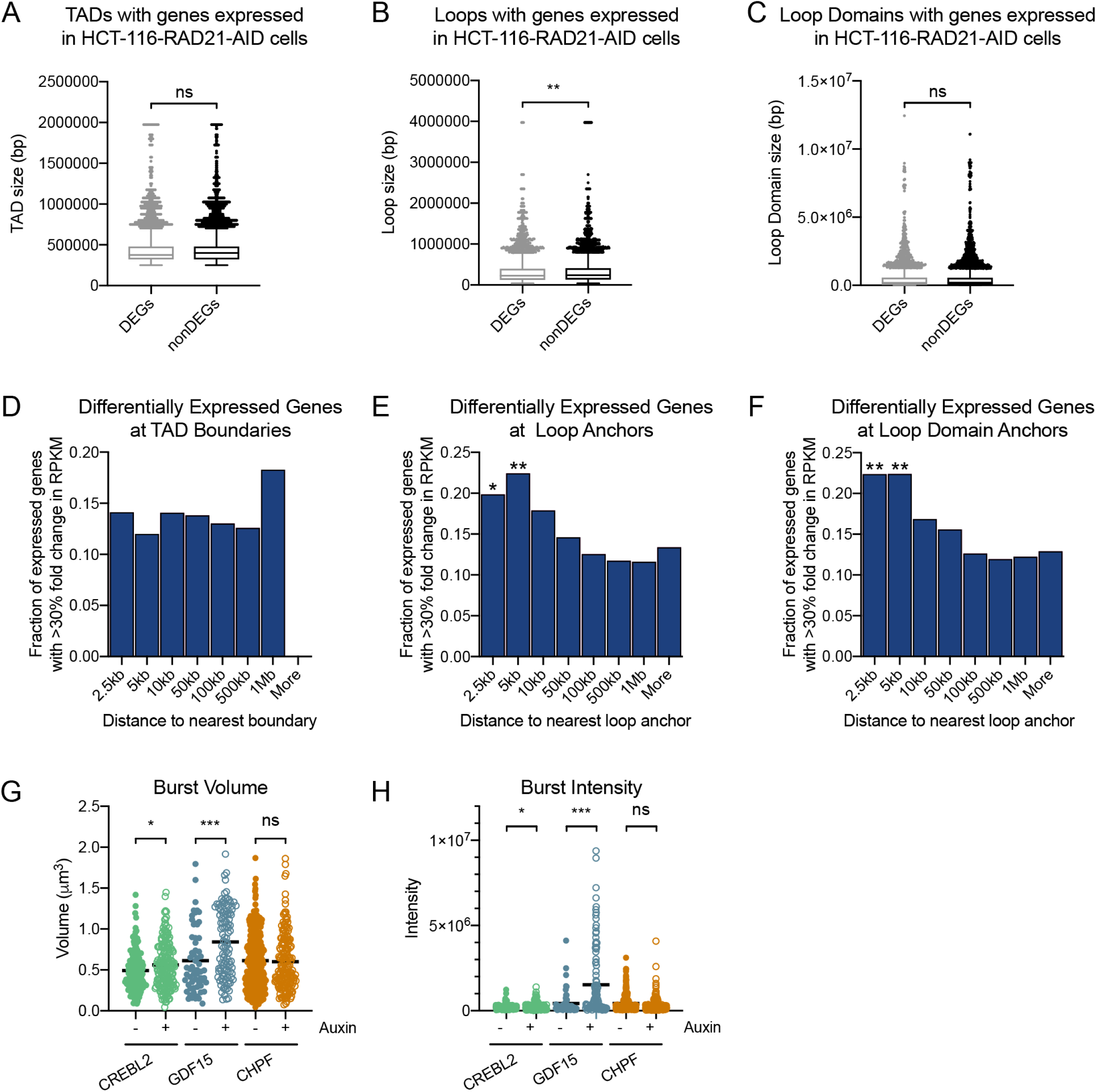
Additional information related to Figures 4 through 6. (A) Distribution of TAD sizes that harbor either a differentially expressed or non-differentially expressed gene in the HCT-116-RAD21-AID cell line. (B) Distribution of loop sizes that harbor either a differentially expressed or non-differentially expressed gene in the HCT-116-RAD21-AID cell line. (C) Distribution of loop domain sizes that harbor either a differentially expressed or non-differentially expressed gene in the HCT-116-RAD21-AID cell line. (D) Bar graphs of the percentage of genes at each binned distance from the TAD boundary that are differentially expressed by >30% fold change upon auxin treatment in the HCT116-RAD21-AID cells. Enrichment was not significant by Fisher’s exact test. (E) Bar graphs of the percentage of genes at each binned distance from the loop anchor that are differentially expressed by >30% fold change upon auxin treatment in the HCT116-RAD21-AID cells. p<0.005, Fisher’s exact test (enrichment within 2.5kb of loop anchor). (F) Bar graphs of the percentage of genes at each binned distance from the loop domain anchor that are differentially expressed by >30% fold change upon auxin treatment in the HCT116-RAD21-AID cells. p<0.005, Fisher’s exact test (enrichment within 2.5kb of loop anchor). (G) Volume of RNA FISH signal per transcriptional burst in HCT116-RAD21-AID cells before (-) and after (+) 6 hours of auxin treatment. * p = 0.018, *** p < 0.001, two-tailed Mann Whitney test. (H) Intensity of RNA FISH signal per transcriptional burst in HCT116-RAD21-AID cells before (-) and after (+) 6 hours of auxin treatment. * p = 0.039, *** p < 0.001, two-tailed Mann Whitney test.

**Table S1:** Architectural design statistics. TopDom algorithm was used to call TADs in Hi-C data from HCT-116-RAD21-AID cells either without (untreated) or following 6 hours of auxin treatment to degrade RAD21 (Rao et al., 2017), as well as from Hi-C from GM12878 (Rao et al., 2014). Table includes the number of domains called and average size of domain. Loop, loop domain, and domain calls were previously published (Rao et al., 2014; Rao et al., 2017) for the cell lines.

| TAD Statistics | HCT-116-RAD21-AID |  |  | GM12878 |  |
| --- | --- | --- | --- | --- | --- |
|  | - Auxin | + Auxin |  |  |  |
|  |  |  | Source |  | Source |
| Number of TADs called | 4429 | 3706 | TopDom | 5056 | TopDom |
| Average size (kb) | 494.18 | 490.14 |  | 494.17 |  |
| Median size (kb) | 425 | 400 |  | 425 |  |
| Number of Loops called | 3170 | 81 | Rao et al.,<br>2017, Cell | 9448 | Rao et al.,<br>2014, Cell |
| Average size (kb) | 364.93 | - |  | 1173.83 |  |
| Median size (kb) | 275 | - |  | 275 |  |
| Number of Loop Domains called | 2104 | - | - | - | - |
| Average size (kb) | 343.93 | 9 (all false positives) | - | - | - |
| Median size (kb) | 265 | - | - | - | - |
| Number of Arrowhead Domains called | - | - | - | 9274 | Rao et al.,<br>2014, Cell |
| Average size (kb) | - | - | - | 258.28 |  |
| Median size (kb) | - | - | - | 185 |  |

**Table S2.** Oligopaint design. Genomic coordinates and number of oligos corresponding to Oligopaint probe designs.

| **Chromosome** | **Paint** | **Start** | **Stop** | **# Oligos** |
| --- | --- | --- | --- | --- |
| 1 | I | 36565548 | 36912169 | 1705 |
| 1 | II | 36912169 | 37924330 | 7418 |
| 1 | III | 37924330 | 38484758 | 2696 |
| 1 | IV | 38484758 | 39301084 | 4288 |
| 1 | V | 39301084 | 40265654 | 3124 |
| 12 | I | 11707040 | 12890169 | 3686 |
| 12 | II | 12890169 | 13408925 | 2097 |
| 12 | S1 | 11707040 | 12189144 | 2078 |
| 12 | S2 | 12189144 | 12487501 | 547 |
| 12 | S3 | 12529811 | 12890169 | 983 |
| 12 | BP; CREBL2 | 12764766 | 12798042 | 56 |
| 22 | I | 33414401 | 35627637 | 7242 |
| 22 | II | 35627637 | 36520199 | 3625 |
| 22 | S1 | 35627637 | 36019409 | 2343 |
| 22 | S2 | 36019409 | 36520199 | 1280 |
| 22 | BP; MCM5 | 35796115 | 35820495 | 196 |

**Table S3.**
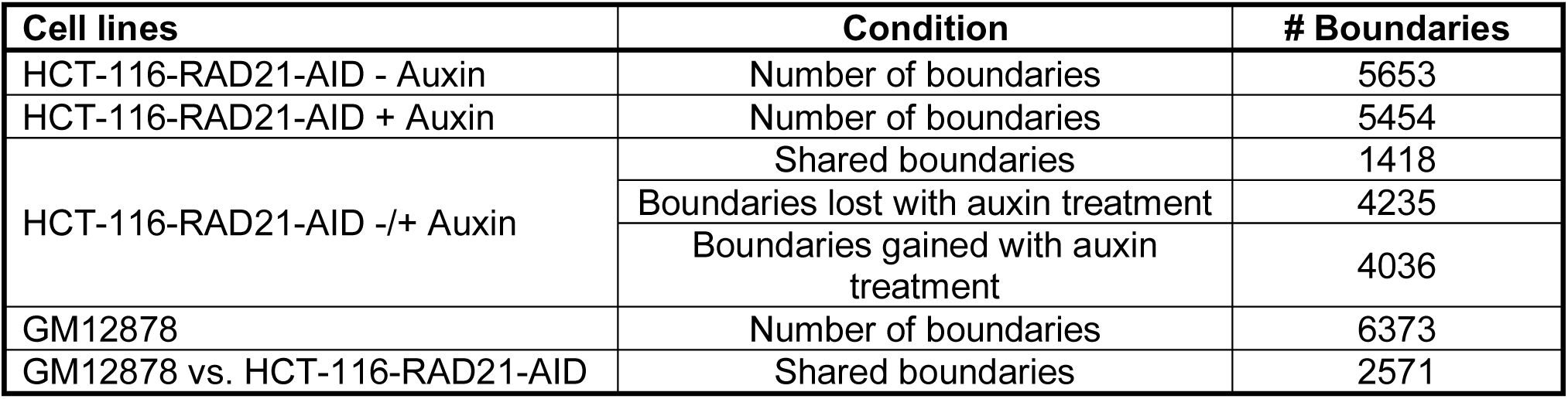
TAD Boundaries. Table detailing the number of shared and boundaries following auxin treatment in the HCT-116-RAD21-AID cell line.

| Cell lines | Condition | # Boundaries |
| --- | --- | --- |
| HCT-116-RAD21-AID - Auxin | Number of boundaries | 5653 |
| HCT-116-RAD21-AID + Auxin | Number of boundaries | 5454 |
| HCT-116-RAD21-AID +/- Auxin | Shared boundaries | 1418 |
|  | Boundaries lost with auxin treatment | 4235 |
|  | Boundaries gained with auxin treatment | 4036 |
| GM12878 | Number of boundaries | 6373 |
| GM12878 vs. HCT-116-RAD21-AID | Shared boundaries | 2571 |

